# Rapid phase resetting of *Aedes aegypti* circadian rhythms by transient alterations in light exposure

**DOI:** 10.64898/2026.08.27.747523

**Authors:** Martin Dessart, Sydney Luff, Lily Smith, Hayden Sunman, Clément Vinauger

## Abstract

Circadian clocks enable mosquitoes to anticipate recurring environmental variations and coordinate behaviors critical for survival and disease transmission, such as locomotion, reproduction, host-seeking, and blood-feeding, with times of day when performance is maximal. In *Aedes aegypti*, locomotor activity follows a robust diurnal rhythm shaped by endogenous circadian clocks and environmental cues, among which light has been shown to be the primary source of temporal information. While early studies established the role of light in regulating locomotor activity, behavior, oviposition and pupation, it remains unclear which features of a light cycle drive changes in circadian rhythms. This question is increasingly relevant as *Ae. aegypti* is frequently exposed to artificial and dynamic lighting conditions in urban environments. Here, we investigated how transient changes in light schedules influence circadian rhythms in locomotor activity by systematically manipulating the timing, duration, and direction of light exposure. Using a high-throughput assay, we tested over 1900 individuals, including wild-type and *timeless* knockout mutants, and showed that a single day of altered lighting is sufficient to induce robust phase shifts, with no evidence of masking effects. A 6-hour light pulse was sufficient to re-entrain mosquitoes regardless of the timing of the pulse, and phase shifts were primarily driven by the offset time of the light pulse, indicating that light-offset acts as a major *zeitgeber*. Together, these findings challenge conventional assumptions about the timescale of circadian synchronization and highlight the remarkable plasticity of mosquito behavior in response to anthropogenic light. Eventually, these effects could explain the rapid adaptation of the species to urban environments and have potential consequences for disease transmission dynamics.

**Highlights:**

- A single day of changes in the light schedule is sufficient to reset the phase of *Aedes aegypti*’s circadian clock.
- Circadian phase resetting requires at least four to six hours of light exposure.
- Advancing the circadian phase is easier than delaying it.

**Graphical abstract:** 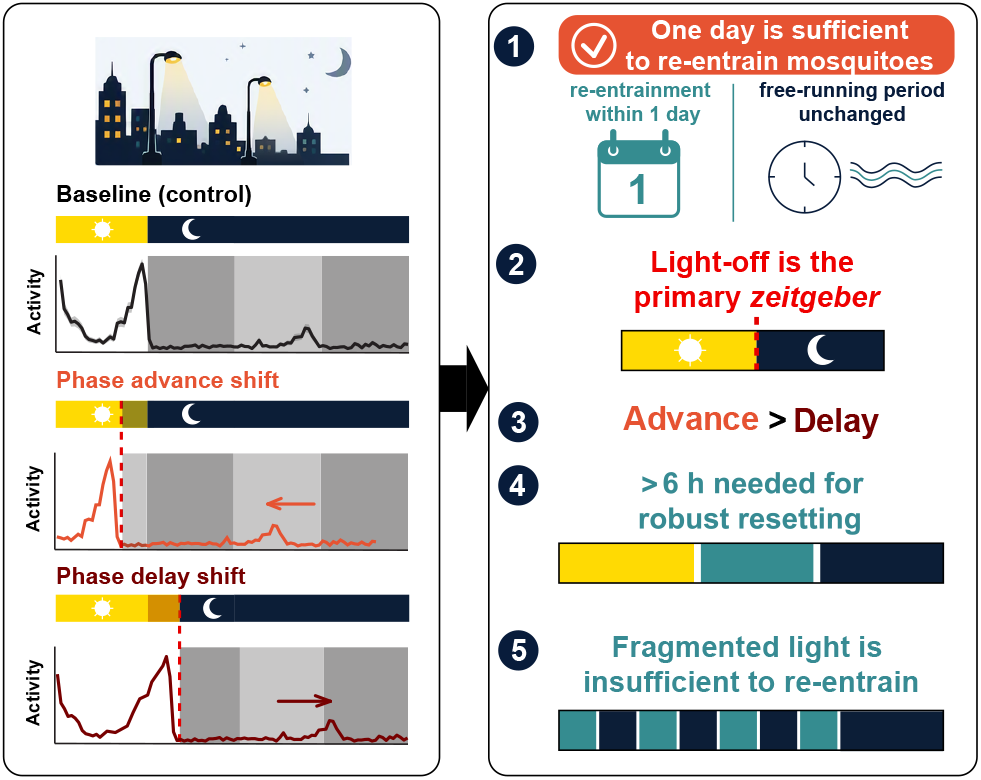

## 1. Introduction

Organisms anticipate predictable environmental changes thanks to internal time-keeping mechanisms, their circadian clocks, which coordinate behavior and physiology with the daily cycles of day and night. These endogenous timing systems regulate a wide range of processes, including locomotion, feeding, reproduction, metabolism, sensory responsiveness, and even insects’ resistance to insecticides (1–9). To remain temporally aligned with local environmental conditions, circadian clocks are continuously synchronized, *i*.*e*., entrained, by external time cues, or *zeitgebers*, among which light is generally the most influential (10–12).

In mosquitoes, circadian regulation plays a central role in behaviors that determine survival and reproductive success (13–16). Flight activity, mating, host-seeking, blood-feeding, and oviposition all occur at specific times of the day and are controlled, at least in part, by endogenous circadian mechanisms (8, 17–22). Among disease-vector mosquitoes, *Aedes aegypti* exhibits a characteristic diurnal activity pattern with a dusk activity peak that persists under constant environmental conditions, demonstrating regulation by an internal circadian clock (18). The timing of these behavioral rhythms has important ecological and epidemiological consequences because it determines when mosquitoes will encounter hosts, mates, suitable environmental conditions, and, ultimately, influences disease transmission (23).

Because light is the primary environmental cue driving circadian entrainment, considerable effort has been devoted to understanding how mosquito clocks respond to changes in illumination. Early studies examined circadian timing in oviposition and pupation, and revealed seasonal adaptations associated with dormancy and latitudinal variation (24– 26); reviewed in (27). In terms of flight activity, pioneering work established that locomotor rhythms in mosquitoes are strongly shaped by the photoperiod and demonstrated that *Ae. aegypti* maintains robust free-running rhythms under constant darkness (18, 28). Later studies characterized phase-response relationships to brief light pulses, providing key insights into how light can advance or delay circadian activity in mosquitoes and potentially influence their vectorial capacity (29–31). Together, these studies used a wide range of experimental standards and methodologies over the last seven decades to establish the role of mosquito circadian clocks and their responsiveness to photic input.

However, the mechanisms by which mosquitoes extract temporal information from complex light environments remain incompletely understood (32). In particular, it is unclear which features of a light cycle are most important in determining the circadian phase. Circadian systems may rely on the timing of dawn, the timing of dusk, day length, previous entrainment, or combinations of these signals to establish temporal alignment with the environment. Distinguishing among these possibilities is critical for understanding how mosquito clocks synchronize to changing photoperiods and how they respond to transient perturbations in light exposure. In *Drosophila melanogaster*, these features have been systematically dissected to reveal how circadian rhythms are generated and synchronized with environmental cues (33, 34), but this remains to be established in mosquitoes.

This question has become increasingly relevant as mosquitoes are now frequently exposed to lighting environments that differ substantially from natural solar cycles (35). With human populations and mosquito vectors becoming more urban and increasingly embedded in human-modified environments, artificial light at night, indoor illumination, and abrupt changes in light exposure are now ubiquitous and can alter the timing, duration, and intensity of photic signals that mosquitoes encounter (36). Yet, it remains unclear how the circadian system interprets such perturbations and whether specific components of the light cycle exert disproportionate control over behavioral timing.

Here, we investigated how *Ae. aegypti*’s circadian rhythms respond to transient alterations in light schedules. By systematically manipulating the timing and duration of light exposure and measuring subsequent locomotor activity under constant-darkness conditions, we tested which temporal features of photic stimuli are most influential in setting the circadian phase. We show that a single day of altered lighting is sufficient to produce robust phase shifts in behavioral rhythms and that these shifts are determined primarily by the timing of light offset and gated by the duration of light exposure. These findings reveal a remarkable capacity for rapid circadian re-entrainment and identify dusk-associated cues (light-offset timing) as an important determinant of circadian timing in *Ae. aegypti*.

## 2. Methods

### 2.1. Mosquito rearing and maintenance

*Aedes aegypti* mosquitoes (Rockfeller F25, MR4-735) were maintained in a climatic chamber at 25 ± 1 C, 60 ± 10% relative humidity (RH) under a 12-12h light-dark cycle with 4 am lighton and 4 pm light-off. Larvae were raised in a 26 x 35 x 4 cm tray filled with 1 cm of deionized water, and fed daily with a Hikari Tropic First Bites (Petco, San Diego, CA, USA). Pupae were isolated on the day of pupation and placed into mosquito breeding containers (BioQuip, Rancho Dominguez, CA, USA). Mosquitoes emerged in the breeding containers and had unrestricted access to 10 % sucrose solution.

### 2.2. Activity monitoring

Mosquito activity was measured using LAM25 actometers (TriKinetics, Waltham, MA, USA). 1-2-day-old mosquitoes were briefly cooled and placed into a transparent Pyrex glass tube with a mesh covering one end and a custom-made 3D-printed cap with a cotton plug soaked in 10% sucrose reservoir on the other end. These tubes were then placed and centered in the actometer so that the infrared beams bisected each tube at its midpoint. Activity counts were recorded as infrared beam breaks as individuals crossed the middle line of the tube. Counts were binned at 1-min intervals. Each actometer was placed in a dark 20-gallon storage bin equipped with a LED light bulb, and transferred to a climatic chamber (model I-36VL, Percival Scientific, Perry, IA, USA, and model 3721 Precision, Thermo Fischer Scientific, MA, USA), maintaining a temperature of 25 ± 1 C and 40 ± 10% RH throughout the experiments. For each actometer, the light-dark cycle was generated using digital timers (model 33860, MyTouchSmart, SunSmart), controlling 800 Lumen light-emitting diodes (LED) (Philips, Amsterdam, The Netherlands). Shifts were performed by manually adjusting the timer’s onset and offset. Due to the iterative nature of the experimental design (*i*.*e*., intermediate results informed subsequent treatment selection), treatments were applied in a pseudo-randomized sequence rather than a fixed or fully randomized order. This approach limited confounding effects of treatment and temporal variation, such as phenological changes, and was validated with additive generalized mixed-effect models, showing that seasonality and replicate identity did not affect the data (see section 2.4 for details). Mosquitoes were left undisturbed for 8-10 days before being examined for survival. Dead individuals were removed from the data (n=327 / 1984, 16.48% of tested individuals), as well as mosquitoes whose activity exceeded 60 beam breaks per hour for one hour, because such sustained activity strongly suggests that the individual was blocked in the center of the beam rather than exhibiting normal locomotor behavior (n=47 / 1984, 2.37% of the dataset). In total, 1610 mosquitoes from 62 independent actometers were retained for the analysis.

### 2.3. Assessing circadian synchronization in mosquitoes

We monitored mosquito locomotor activity using actometers under controlled light-dark (LD) conditions, and applied a series of controlled temporal shifts (Fig. 1). After either 1 or 4 days of entrainment to a 12:12 LD cycle, mosquitoes were subjected to a phase shift (advance or delay). Individual activity was then recorded for three consecutive days in constant darkness (DD) to quantify their endogenous circadian activity. To distinguish acute phase shift responses from stabilized circadian resynchronization, we compared the time of peak activity on the first day in DD with the average peak time across three days in DD. Given that in *Ae. aegypti*, the morning activity peak is absent under DD conditions, we focused our analysis on the dusk activity peak.

**Fig 1.**
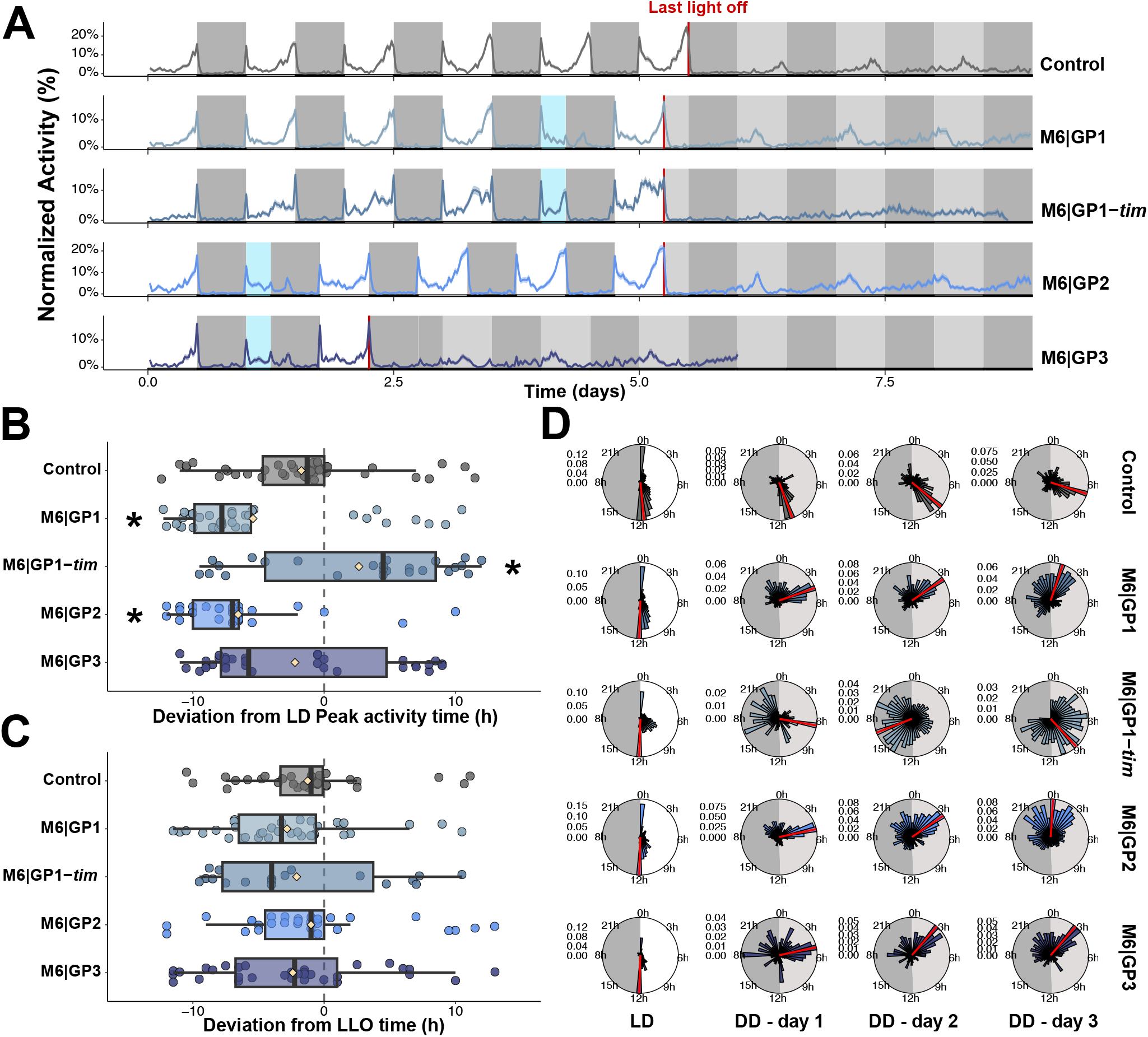
Mosquitoes can rapidly re-entrain to a 6-h phase advance shift with a 24-h recovery period contingent on a functional clock and sufficient prior entrainment. (A) Population activity profiles. Locomotor activity is shown as Williams’ mean over time, with 1 bin per minute. Shaded ribbon represents the mean ± standard error of the mean (SEM). Mosquitoes were exposed to an 6-h advanced shift (lights off at ZT6 instead of ZT12) followed by either one or four post-shift entrainment days. M6|GP1 (n = 74): four days of LD entrainment, one day shift and one day post-shift entrainment; M6|GP1-*tim* (n = 76): *timeless* loss-of-function mutants subjected to the same treatment as the M6|GP1 group; M6|GP2 (n = 58): one day of LD entrainment, one day shift and four days of post-shift entrainment; M6|GP3 (n = 64): one day of LD entrainment, one day shift and one day of post shift entrainment at an earlier age. Background shading indicates the imposed lighting schedule: white and black regions represent the light and dark phases of the LD cycle; blue regions indicate the experimental light-treatment period (*i*.*e*., shift); and grey/black regions indicate constant-darkness (DD) interval and projected subjective light/dark phases. Red vertical lines mark the last lights-off time (LLO). **(B-C)** Phase deviation of activity peaks timing relative to **(B)** baseline peak activity time under LD and **(C)** the time of last light-off event. Dashed vertical line denotes no shift relative to the peak timing under the preceding LD cycle or relative to the LLO, respectively. Negative and positive values indicate activity peaks occurring earlier and later, respectively. Boxes show the interquartile range and median, whiskers indicate the distribution spread, points represent individual observations, and beige diamond-shaped symbols indicate group means. **(D)** Left to right: Rose diagrams showing the mean population activity by day calculated in 30-min bins across 24 h, for: baseline LD, and days 1, 2 and 3 under DD (DD1-DD3). Under LD conditions, the shaded sector denotes the dark phase. Under DD conditions, the light- and dark-grey sectors indicate the subjective day and subjective night, respectively. The red radial line marks the time bin containing the activity peak. Each panel is normalized to its own maximum activity.

### 2.4. Quantification and statistical analysis

All analyses were performed in *R* version 4.5.0 (2025-04-11), using the *Rethomics* framework (37). Raw activity counts were grouped by individual, log-transformed and averaged using Williams’ mean using the *WPR* function from the *TTR* package (38), then re-centered by subtracting 0.5 to yield a normalized activity score. Rose diagrams were plotted using the *circular* package (39). For each individual, the free-running period was computed using an autocorrelation-based periodogram that determines a significance threshold based on a null distribution (*p* = 0.05). Peaks above this threshold were considered significant. Under constant darkness, one peak per 24-h period was counted, and individuals with at least one significant peak were classified as rhythmic. A total of n= 1400 / 1610 (86.96%) mosquitoes were considered rhythmic, and the remaining n= 210 / 1610 (13.04%) mosquitoes were excluded from subsequent phase deviation analyses.

Williams’ mean activity was averaged by 30-minute bins, and mean activity levels were calculated for each individual. To quantify the change in the timing of the activity peak, we identified the time bin corresponding to peak activity for each individual and each 24-h period. The peak activity times were then averaged per treatment using a custom-written circular mean function that converted time values from hours to radians and back to hours.

From these peak activity values, two complementary metrics were defined to calculate changes in activity: *(i)* the phase deviation between the time of peak activity in DD relative to each individual’s peak activity time under LD cycle, reflecting the effect of the light-induced phase shift on the endogenous circadian clock, and *(ii)* the phase deviation between the time of peak activity in DD and the time of the last lights-offset (LLO), reflecting the degree of alignment to the new, shifted, light schedule.

For each metric, and over the first day in DD or across the three DD days, we fitted a generalized linear mixed-effects model using the *glmmTMB* package (40), with phase deviation as the response variable and the experimental treatment as the explanatory factor. Random intercepts included actometer identity and tube positioning within the actometer to account for non-independence. As experiments were conducted over a 20-month period, we also controlled for potential seasonal and replicate effects by adding the actometer setup date x treatment replicate as random effects. Deviations were compared with the control group using post hoc Dunnett-adjusted contrasts. No significant effects of date or actometer identity, as well as interactions between experience and date were detected (*p* > 0.1).

When appropriate, differences in dispersion between groups were tested using the Walraff Test of angular distances, using *wallraff*.*test* function from the *circular* package (39). For each individual, the mean activity level in DD was calculated by averaging the Williams’ mean activity per hour and normalized by subtracting the corresponding mean activity in LD. Mean activity was analyzed using generalized linear effects models using the *glmmTMB* package followed by Tukey-adjusted post-hoc comparisons against the control. Differences in the proportion of rhythmic individuals between groups were evaluated using a generalized linear model with a binomial distribution using the *glmmTMB* package with Tukey-adjusted post-hoc comparisons. Finally, the free-running period was compared across treatments using a generalized linear model using the *glmmTMB* package. No significant differences were observed between treatments (*p* = 0.88).

## 3. Results

### 3.1. A single day following a 6-h advance shift is sufficient to robustly phase-shift mosquitoes

As a first challenge to the circadian clock, we introduced a temporal shift of 6 h after four days of entrainment in 12:12 L:D, followed by one day of entrainment aligned with the shifted (advanced) LD cycle (*i*.*e*., lights on at ZT18 - lights off at ZT6 relative to the original ZT0-ZT12 cycle, Fig. 1A,B, S1,S2). In constant darkness (DD), mosquitoes showed a significant phase shift compared to the control (*p* < 0.0001, Control: −1.72 [95% CI −2.95; −0.48], M6|GP1: −7.95 [95% CI −9.01; −6.88]), consistent with a 6 h advance (Fig. 1, C). Moreover, no differences were observed between their deviation from the time of last lights-off (LLO) time when compared to that of the control group on both the first day and within three days since LLO (*p* > 0.56, (Fig. 1D, S1F).

Using a *timeless* knockout line (41), we performed a similar 6-h advance shift on clock-deficient mosquitoes to confirm the endogenous nature of the rhythm. *timeless* is a core clock gene that encodes the TIMELESS (TIM) protein, which partners with PERIOD (PER) to form a complex that inhibits the transcriptional factors CLOCK (CLK) and CYCLE (CYC). In mosquitoes, light-activated CRYPTOCHROME1 degrades TIM, which in turn enables the chelation of CLK:CYC by the CRY:PER dimer and allows a new cycle. In consequence, *timeless* knockout mutants can still show rhythmic activity under an LD cycle but cannot sustain a self-maintained endogenous circadian rhythm under constant conditions (41). Following the 6-h advance shift, *timeless* mutants showed a significantly reduced rhythmicity in DD compared to control mosquitoes (binomial GLM, *n*_control_ = 61, *n*_timeless_ = 76, *p* < 0.001), with only 65% of *timeless* mutants classified as rhythmic and 92% for the Control (S1C), confirming that phase shifts in wild type individuals are mediated by the endogenous circadian clock, and that a functional clock is necessary to sustain rhythmic activity after a phase shift.

Next, we tested the effects of the duration of post-shift entrainment (1 day versus 4 days; Fig. 1) and controlled for age by comparing shifts occurring on days 1 or 4 after the start of the assay. With four days of post-shift entrainment (M6|GP2), we observed a similar pattern to one day of postshift entrainment occurring on day 4 (M6|GP1), with a significant phase shift (*p* < 0.0001, M6|GP2: −7.74 [95% CI −8.91; −6.57], Fig. 1). In contrast, when younger mosquitoes were provided with only one day of post-shift entrainment (M6|GP3), they slowly aligned their peak activity time with the new LD cycle but did not show a clear phase shift on the first day under DD (*p* = 0.54, M6|GP3: −4.05 [95% CI −6.93; −1.16], Fig. 1). Although the mean of the deviations were not different from the Control, the M6|GP3 treatment showed a significantly greater dispersion (Wallraff’s Test: *p* < 0.01), indicating that some individuals were shifted while others remained on the initial cycle (Fig. 1).

While the time of peak activity under DD conditions was significantly affected by the experimental treatments during the 1st day and the average of 3 days in DD (*p* < 0.0001), all groups showed a drift in peak activity timing reflecting a freerunning period shorter than 24 h. The experimental treatments did not significantly impact the length of the free running period (*p* = 0.88). Likewise, estimated marginal means for the LD–DD difference in activity ranged from −2.00% to 0.43% across conditions, indicating a small, broadly consistent reduction in activity from LD to DD (Fig. S1 to Fig. S12). Significant differences with control were observed only for M6|GP2 and M6|GP3 (*p* < 0.02), consistent with 4 days post-entrainment and younger mosquitoes respectively.

### 3.2. Advance shifts can be reverted, affect males as well, and do not require post-shift entrainment

We next wondered whether the effects of an advance shift could be reverted by post-shift entrainment to the original LD cycle. To test this, we provided shifted mosquitoes with a 24 h postshift entrainment (*i*.*e*., a full 12:12 LD 24 h day) aligned with the original LD cycle (ZT0-ZT12) (Fig. 2 M6|GP1-NS). In this case, mosquitoes rapidly reverted to the initial phase, showing no shift in DD and no significant deviation from the original cycle (deviation in DD: *p* = 0.52, M6|GP1-NS: −3.25 [95% CI −4.92; −1.58]; deviation since LLO *p* = 0.99, M6|GP1-NS: −1.63 [95% CI −3.35; −0.09]).

**Fig 2.**
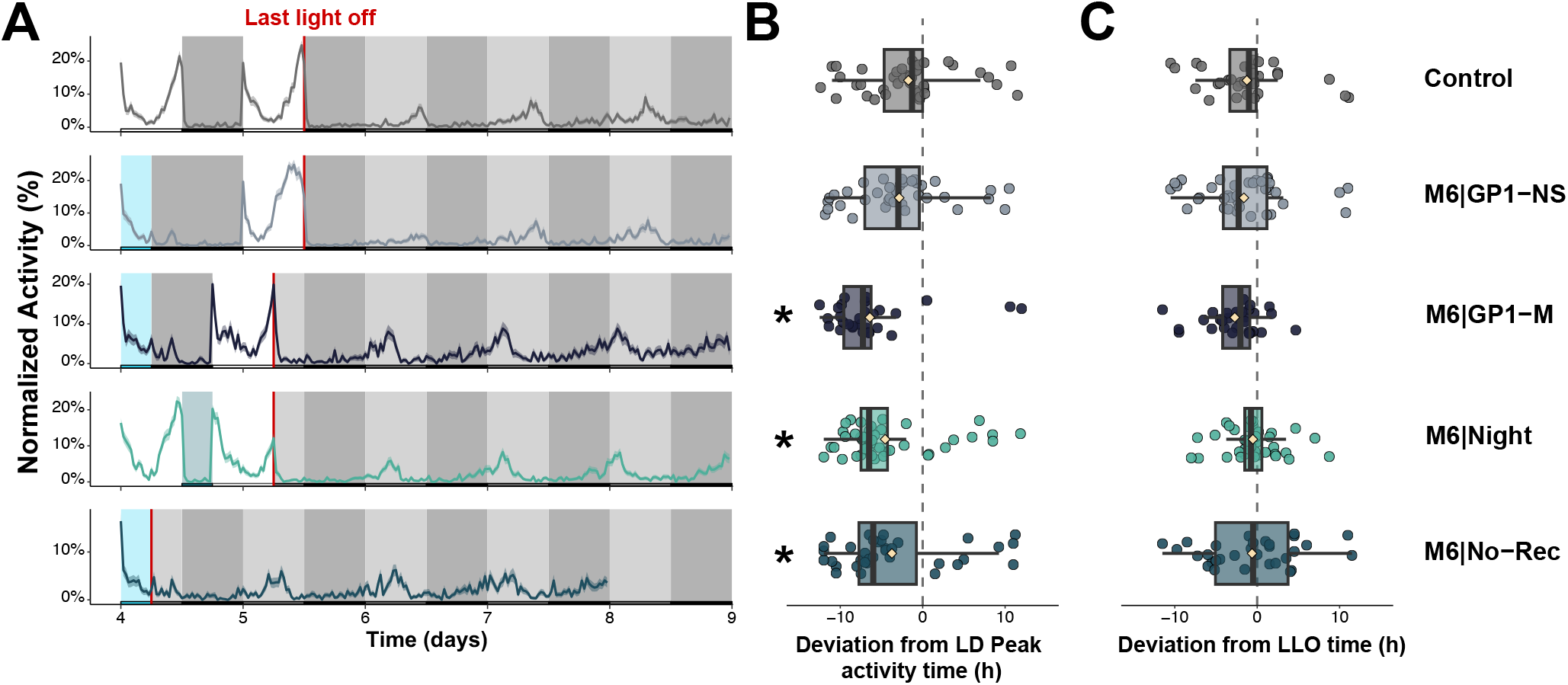
Re-entrainment to a 6-h advance shift occurs in males, is independent of the phase and doesn’t require post-shift entrainment, but can be canceled by a single recovery day. **(A)** Population activity profiles of: mosquitoes exposed to a 6-h advance shift under a 24-h recovery period synchronized to the original LD cycle (M6|GP1-NS: n = 57); males exposed to the 6-h advance-shift (M6|Male: n = 48), females exposed to the shift during the scotophase (M6|Night: n = 67), or following immediate transfer to DD without post-shift entrainment (M6|No-Rec: n = 57). For visual and direct comparison across treatments, the Control group is replotted from Fig. 1. Activity is shown as Williams’ mean and shaded ribbon represents the mean ± standard error of the mean (SEM). Background shading indicates the imposed lighting schedule: white and black regions represent the light and dark phases of the LD cycle, blue regions indicate the experimental light-treatment period; and grey and black regions indicate the subsequent constant-darkness interval and its projected subjective day and night, respectively. Red vertical lines indicate the LLO time. **(B-C)** Phase deviation of activity peaks timing relative to **(B)** baseline activity peak time and **(C)** the LLO event. Each point represents an individual’s activity peak phase. Dashed vertical lines denote no shift relative to the peak phase under the preceding LD cycle, or to the LLO time, respectively. Boxes show the interquartile range and median, whiskers indicate the distribution spread, points represent individual observations, and beige diamond-shaped symbols indicate group means.

We also compared males and females following a 6 h phase advance shift followed by one day of post-shift entrainment and found that males shifted as well as females (*p* < 0.0001, M6|Male: −7.68 [95% CI −8.61; −6.76], Fig. 2, S3, S4). Given the absence of sex-specific differences and the epidemiological importance of females, subsequent experiments focused on females only.

We then applied the 6 h advance shift during the scotophase instead of the photophase by shortening the night (from ZT12 to ZT18) instead of the day. In this scenario, the first day in DD was clearly shifted (*p* < 0.0001, M6|Night: −6.52 [95% CI −7.33; −5.70], Fig. 1). Similarly, no significant deviation to the LLO was detected in night-shifted mosquitoes for the first day and across three days after LLO (*p* > 0.90, Fig. 1, S3F).

Because mosquitoes readily entrained to the shifted LD cycle across all the experiments presented so far, we tested whether mosquitoes could entrain to a 6 h advance shift, even without a post-shift entrainment day. To do this, the shift was immediately followed by DD conditions (Fig. 2). Under these conditions, a significant phase shift was observed after 1 day in DD (*p* < 0.0001, M6|No-Rec: −5.99 [95% CI −7.66; −4.33], Fig. 2), was sustained across 3 days in DD (*p* < 0.0001, Fig. S3, S4), and no significant deviation from the LLO time was detected across three DD days (Compared to control conditions: *p* > 0.89, Fig. 1, S3F).

### 3.3. Delay shifts entrain mosquitoes but with reduced efficiency compared to advance shifts

We next examined the impact of the shift direction by applying a 6 h delay shift (*i*.*e*., making the day or the night longer). Following 6 h phase delay applied during the photophase and one day of post-shift entrainment (P6|GP1), mosquitoes showed a significant phase shift in DD (*p* < 0.001, first day: 2.92 [95% CI 1.60; 4.24], three-day average: 1.87 [0.90; 2.86], Fig. 3, S5,S6) and no significant deviation from LLO time across three DD days (*p* > 0.22, Fig. 3, S5F).

**Fig 3.**
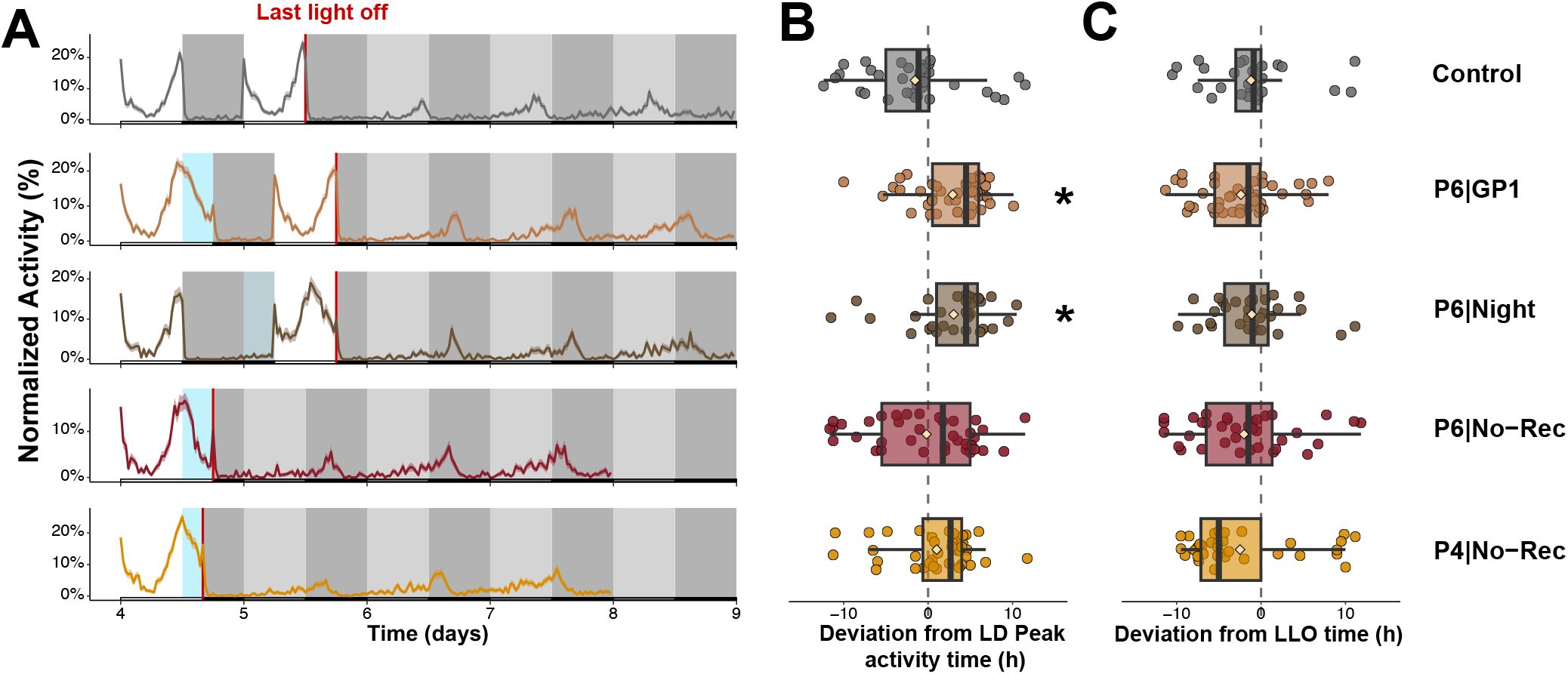
Delay shifts entrain mosquito activity, but require post-shift entrainment. **(A)** Population activity profiles of mosquitoes exposed to: a 6-h delay shift during the photophase (P6|GP1: n = 62), during the scotophase (P6|Night: n = 52), or during the photophase and without post-shift entrainment (P6|No-Rec: n = 59). P4|No-Rec (n = 57) mosquitoes were exposed to a 4-h delay shift without post-shift entrainment. For visual and direct comparison across treatments, the Control group is replotted from Fig. 1. Activity is shown as Williams’ mean, and shaded ribbon represents the mean ± standard error of the mean (SEM). Background shading indicates the lighting schedule: white and black regions represent the light and dark phases of the LD cycle, blue regions indicate the experimental light-treatment period, and grey and black regions indicate the subsequent DD and its projected subjective day and night, respectively. Red vertical lines indicate the LLO time. **(B-C)** Phase deviation of activity peaks timing relative to **(B)** basline peak activity time and **(C)** the LLO time. Each point represents an individual’s activity peak time. Dashed vertical lines denote no shift relative to the baseline peak time or LLO time, respectively. Boxes show the interquartile range and median, whiskers indicate the distribution spread, points represent individual observations, and beige diamond-shaped symbols indicate group means.

When the shift was applied during the scotophase (ZT12-ZT24; P6|Night), similar effects were found. The first day in DD as well as three day average were significantly shifted (*p* < 0.001, first day: 3.033 [95% CI 1.29; 4.78], three-day average: 1.45 [0.45; 2.45], Fig. 3, S5), and no deviation to the LLO time was detected both for the first DD day and across the three DD days (*i*.*e*., no difference from the control’s deviation were found: *p* > 0.6, Fig. 3, S5F).

Conversely to advance shifts, mosquitoes did not exhibit significant changes of peak activity time in the absence of post-shift entrainment for both the first day and the three DD days average (P6|No-Rec: first day: *p* = 0.62, −0.19 [95% CI −2.26; 1.88], three-day average: *p* = 0.74, - 1.71 [−2.77; −0.66], 3, S5E), indicating that advances were more effective than delays in driving circadian adjustment. For the first day in DD, the dispersion of the data was significantly greater than the Control (Wallraff’s Test: *p* < 0.02), indicating that some individuals could re-entrain to the new cycle while others remained on the initial one (Fig. 3, S5, S6).

We next assessed whether a shorter shift would produce a greater effect (*i*.*e*., by representing a smaller challenge to the circadian clock) by applying a 4-hour delay shift without post-shift entrainment. Under this condition, mosquitoes did not shift the time of their peak activity either (P4|No-Rec: first day: *p* = 0.08, 1.03 [95% CI −0.57; 2.63], three-day average: *p* = 0.99, −2.21 [−3.21; −1.21], (Fig. 3, S5), while the dispersion remained similar to the Control (Wallraff’s Test: *p* = 0.75, S5, S6).

### 3.4. When the lights-off time is aligned with the original LD cycle, the duration of the shift has no effects

To further investigate the effect of the shift duration, we applied a series of light stimuli from 1 to 8 h in duration, with their end aligned with the lights-off time (ZT12) under the initial LD cycle (Fig. 4A). Regardless of the light stimulus duration, no immediate phase shift was observed after one day in DD (*p* > 0.2 for all conditions, Fig. 4). However, 6 and 8 h of light induced a significant shift across three days in DD, suggesting either a shortening of the free-running period or a larger amount of inter individual variation on the first DD day (OF|6: *p* < 0.01, −4.29 [95% CI −5.26; −3.33], OF|8:*p* < 0.01, −4.27 [95% CI −5.10; −3.45], Fig. S7,S8E). None of these groups showed a deviation from the LLO time that was different from the control (*p* > 0.1, Fig. 4, S7F). This indicates that either prior entrainment to the LD cycle predominated over the shifted light exposure or that the lights-off event signals the shifted phase to the circadian clock.

**Fig 4.**
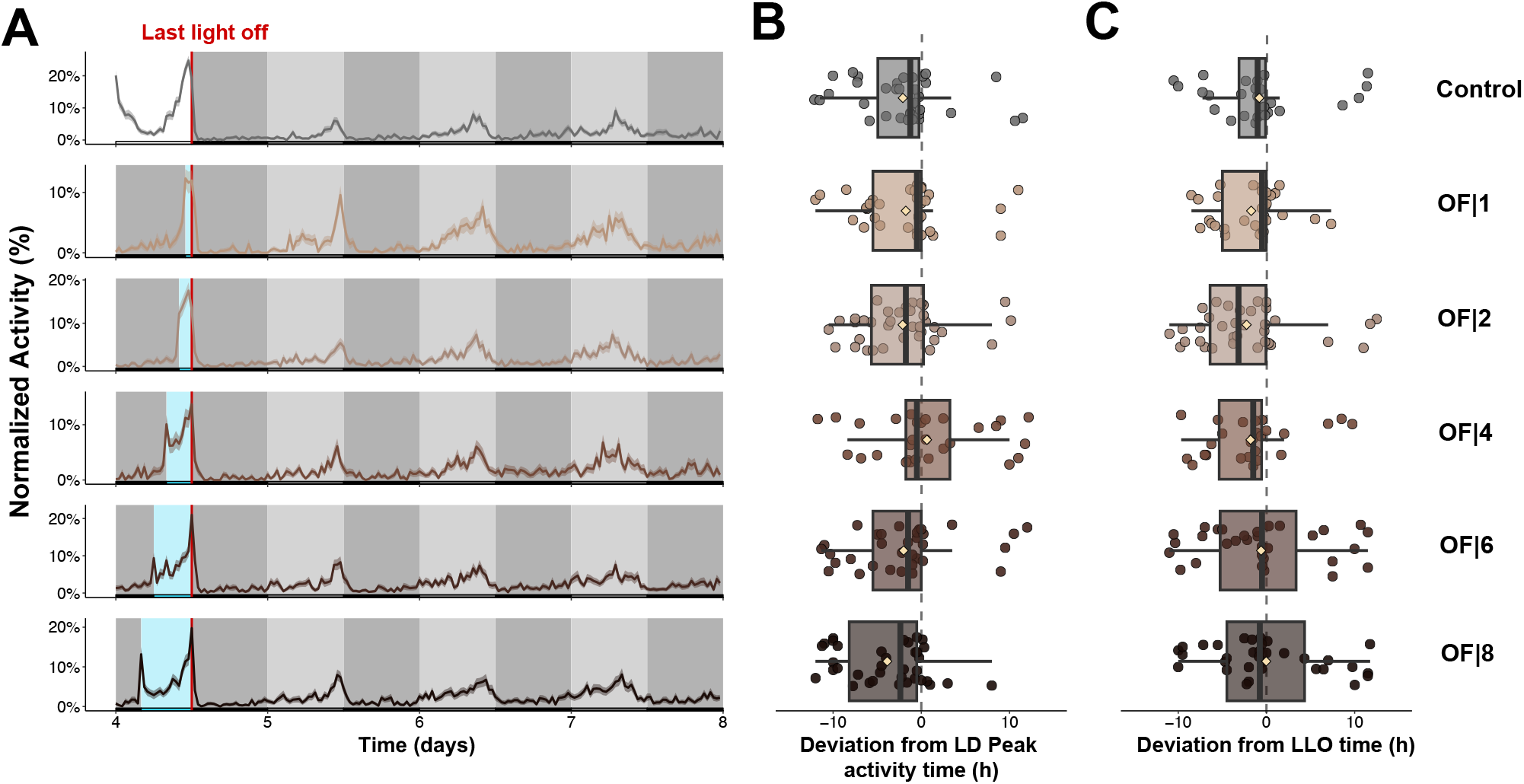
Shifts whose lights-off are aligned with the baseline LD cycle have no effect on the peak activity time, unless long duration stimuli are introduced. **(A)** Population activity profiles of mosquitoes exposed to light duration ranging from 1 to 8 h (OF|1: n =54, OF|2: n = 56, OF|4: n = 54, OF|6: n = 55, OF|8: n = 56), all ending at the original lights-off time (ZT12). For visual and direct comparison across treatments, the Control group is replotted from Fig. 1. Activity is shown as Williams’ mean and shaded ribbon represents the mean ± standard error of the mean (SEM). Background shading indicates the lighting schedule: white and black regions represent the light and dark phases of the LD cycle, blue regions indicate the experimental light-treatment period; and grey and black regions indicate the subjective day and night under DD conditions. Red vertical lines indicate the LLO time. **(B-C)** Phase deviation of activity peak timing relative to **(B)** baseline peak activity time in LD and **(C)** the LLO time. Each point represents an individual’s peak activity time. Dashed vertical lines denote no shift relative to the baseline peak time or LLO time, respectively. Boxes show the interquartile range and median, whiskers indicate the distribution spread, points represent individual observations, and beige diamond-shaped symbols indicate group means.

### 3.5. Lights-off acts as the primary *Zeitgeber* and phase resetting was consistently observed following six-hour light exposures

To disentangle the effects of the lights-off time and the duration of the light exposure, we subjected mosquitoes to a series of light stimuli aligned with lights onset under LD (ZT0) (Fig. 5). Short-duration stimuli (*i*.*e*., 1 and 2 h) did not lead to any significant shift during the first DD day (*p* > 0.25) but the deviation between their peak activity time under DD and the LLO time was significantly different from the control (*p* < 0.0001, (ON|M1: 7.85 [95% CI 6.36.45; 9.35], ON|M2: 6.39 [95% CI 4.52; 8.27], Fig. 5B). However, longer light exposures (6 and 8 h) significantly shifted the time of peak activity from its baseline under LD (ON|M8: first day: *p* < 0.03; −4.39 [95% CI −5.45; −3.34], three-day average: *p* < 0.0001; −5.31 [95% CI −6.01; −4.60], Fig. 5B, S9E), and no significant deviation to the LLO time was detected (*p* > 0.9 for all conditions, Fig. 5C, S9, S10

Mosquitoes subjected to a 4-h long stimulus did not significantly shift from the baseline peak activity time (*p* = 0.9; −2.88 [95% CI −5.05; −0.72], Fig. 5B), while their peak activity time under DD was not significantly different from the LLO time either (*p* = 0.75; −0.20 [95% CI −2.06; 1.66], Fig. 5C). However, the dispersion of this group was significantly greater than that of the Control (Wallraff’s Test: *p* < 0.01, 5B), suggesting partial shift of the group with some individuals shifting their activity and others not.

**Fig 5.**
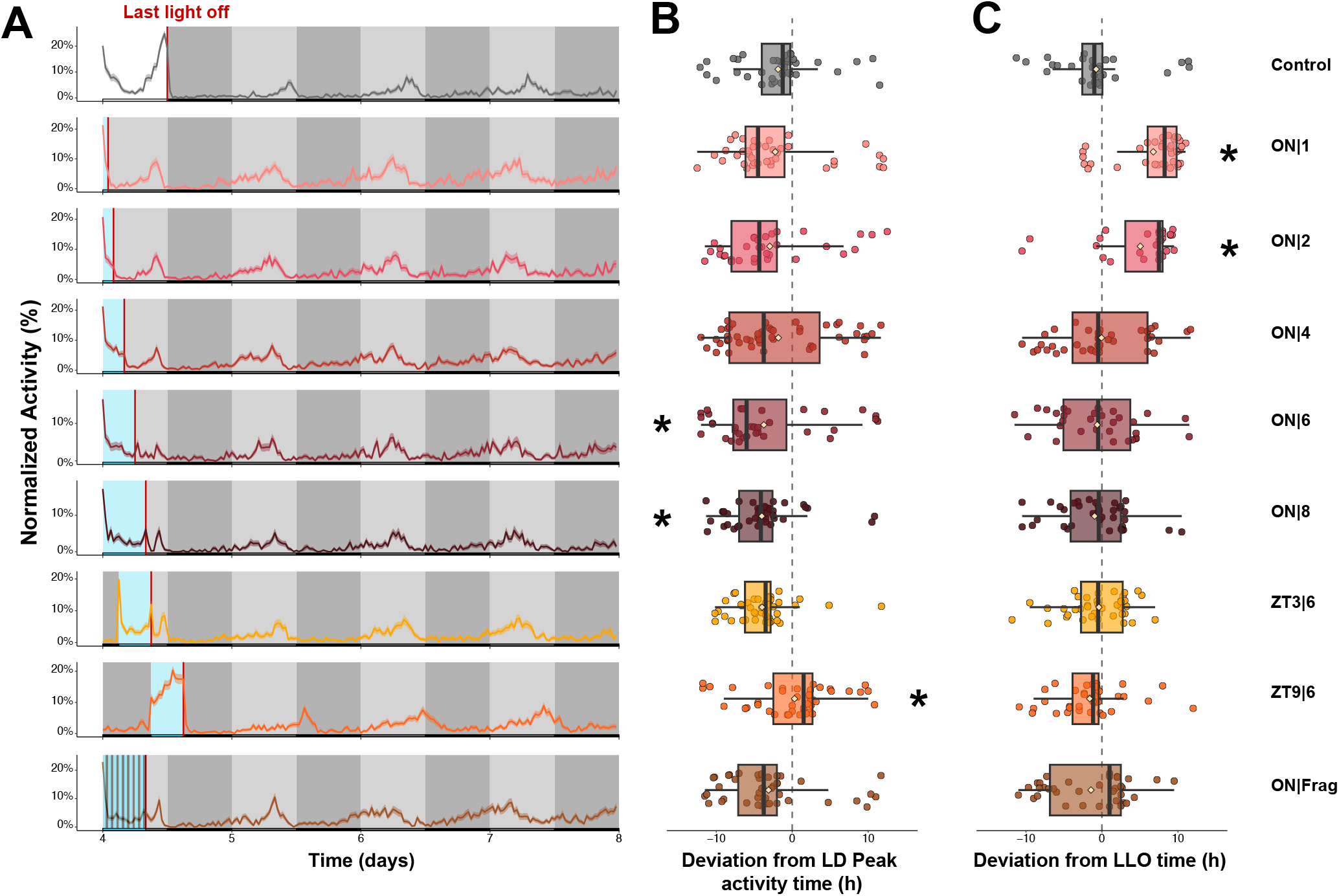
The timing, duration, and continuity of light exposure affect the successful re-entrainment of mosquitoes. **(A)** Population activity profiles of mosquitoes exposed to different light duration and alignments. Light stimuli were either aligned with lights-on (ZT0; ON|1: n = 56, ON|2: n = 50, ON|4: n = 75, ON|6: n = 57, ON|8: n= 56), delivered for 6-h at different circadian times (ZT3|6: n = 57 and ZT9|6: n = 75), or fragmented into alternating 30-min light and 30-min dark intervals over 8 h (ON|Frag; total light exposure = 4 h). ON|6 treatment corresponds to M6|No-Rec treatment in (Fig. 2). For visual and direct comparison across treatments, the Control group is replotted from Fig. 1. Activity is shown as Williams’ mean, and shaded ribbon represents the mean ± standard error of the mean. Background shading indicates the light schedule: white and black regions represent the light and dark phases of the LD cycle, blue regions indicate the experimental light-treatment period; and grey and black regions indicate the subjective day and night under DD, respectively. Red vertical lines indicate the LLO time. **(B-C)** Phase deviation of activity peaks relative to **(B)** baseline peak activity time under LD and **(C)** the LLO time. Each point represents an individual’s activity peak time. Dashed vertical line denotes no shift relative to the peak activity time under LD or relative to the LLO time, respectively. Boxes show the interquartile range and median, whiskers indicate the distribution spread, points represent individual observations, and beige diamond-shaped symbols indicate group means.

To decouple the on and off times of the photic stimuli from the peak and troughs of clock genes’ expression cycles, we further tested 6 h phase advance shifts applied at different *Zeitgeber* times (ZT3 and ZT9) that do not correspond to 1*/*2 or 1*/*4 periods of the circadian clock (Fig. 5, S11, S12). Under these conditions, phase deviations in DD depended strongly on the timing of when the light stimuli turned off (ZT0|6: −5.99 [95% CI −7.66; −4.33], ZT3|6: −4.15 [95% CI −5.01; −3.28], ZT6|6: −2.16 [95% CI −4.40; 0.07], ZT9|6: 0.83 [95% CI −0.60, 2.26], Fig. 5, S11E), indicating that the end of the photic stimulus is the most important driver, with the most pronounced shift magnitudes observed when lights-off were the furthest away from the peak time in LD.

While ON|6 was significantly shifted both on the first day and across three days in DD (*p* < 0.001), ZT3|6 did not reach statistical significance on the first DD day after Dunnett’s correction (*p* = 0.0518) and was significantly shifted across the three DD days (*p* < 0.001, Fig. 5, S11E). ZT6|6 was only shifted when considering the average across the three DD days (*p* < 0.01), whereas ZT9|6 was shifted only on the first DD day (*p* = 0.05, Fig. 5, S11E). None of these groups’ deviation from the LLO time was significantly different from the control (*p* > 0.05, Fig. 5C, S11F).

Finally, we asked whether the amount of light required to entrain mosquitoes needs to be delivered continuously, or whether shorter bouts of light stimuli are sufficient, cumulatively, to produce the same effect. Specifically, we tested the effect of a fragmented light stimulus consisting of alternating 30-minute light and 30-minute dark periods over 8 h (total light exposure = 4 h, ON|Frag; Fig. 5, S11).

Mosquitoes subjected to the fragmented treatment did not recapitulate the effects shown in ON|8 (*i*.*e*., a clear shift of the peak timing in DD). The trend was similar to the ON|4 group, showing a slight tendency to shift their activity peak, but not significantly from Control, both during the first DD day and across the three DD days (first day: *p* = 0.17 - 4.05 [95% CI −5.41; −2.69], three-day average: *p* = 0.06, −4.07 [95% CI −5.18; −2.97], Fig. 5, S11E). No significant deviation from the LLO time was observed (*p* > 0.88, Fig. 5, S11F), but a significant increase in the dispersion of the deviation from LLO time (Wallraff’s Test: *p* < 0.001) was noticed on the first DD day (5C). Moreover, across the three DD days, ON|4 was deviated positively, closer to ON|2, while ON|Frag was deviated in the negative direction (Fig. S9, S11).

## Discussion

In this work, we showed that mosquito circadian clocks respond rapidly to changes in environmental light cues, with no evidence of a delayed masking effect following phase shifts. Overall, robust phase resetting emerged under treatments containing at least six hours of continuous illumination, but the robustness and uniformity of this response depended on the duration of prior LD entrainment, the direction of the shift, the distance from the baseline peak activity time, and the exposure to a post-shift entrainment under LD. Indeed, longer pre-shift entrainment and post-shift re-entrainment led to consistent and population-wide phase shifts that aligned more accurately with the new, shifted cycle. Conversely, shorter pre-shift entrainment produced heterogeneous responses, with partial or heterogeneous shifting across individuals. The time at which the photic stimuli were turned off explained mosquitoes’ ability to shift their peak activity time relative to their baseline. Lights-off that were too close to the baseline activity peak did not induce a significant shift (*e*.*g*., ON|M1), and lights-off that were too far (*e*.*g*., ON|M8) likely challenged the circadian clock to entrain, especially in the absence of post-shift entrainment.

Even without a post-shift entrainment, mosquitoes exposed to a 6-h phase advance exhibited immediate reentrainment of their endogenous clock, an effect primarily driven by the lights-off transition. Importantly, assays with light stimuli aligned with the light onset timing of the baseline photoperiod suggest that a minimum duration of 6 h was required to re-entrain mosquitoes to a new cycle. However, given that shorter duration stimuli also imply a lights-off time that is more distant from the baseline peak activity time, further work will be required to test the re-entrainment to shorter stimuli that ends at Zeitgeber times to which mosquitoes could entrain (*e*.*g*., ON|6).

Although our experiments were not designed to identify the molecular mechanisms underlying phase resetting, the observed patterns suggest several hypotheses. First, the prominent effect of light-offset timing supports a model in which the mosquito circadian clock is particularly sensitive to light-offsets, consistent with earlier observations in *Ae. aegypti* that both lights-on and lights-off can act as phase-setting cues, but that activity timing remains closely associated with the evening transition (18). Our results are broadly consistent with the classical model of circadian photic entrainment established in *Drosophila melanogaster*, in which light induces degradation of TIM and thereby resets the phase of the molecular oscillator. In *Drosophila*, the timing of TIM degradation relative to the ongoing molecular cycle determines whether light produces advances or delays in behavioral rhythms (42). The reduced rhythmicity observed in *timeless* mutants following phase shifts strongly supports the conclusion that phase resetting depends on the molecular circadian clock and is consistent with a conserved TIM-dependent entrainment pathway in mosquitoes. However, our data also indicate that the timing of light offset is a stronger predictor of behavioral phase than light onset, raising the possibility that mosquito clocks integrate photic information over extended periods and use the transition to darkness as a critical reference point for phase resetting. Testing this hypothesis will require direct measurements of TIM and PER dynamics during and after shifted light schedules. Second, the greater effectiveness of phase advances compared with delays may reflect the intrinsic properties of the underlying oscillator. Because the free-running period of *Ae. aegypti* is shorter than 24 h, advances require adjustments in the same direction as the endogenous tendency of the clock to run fast, whereas delays require slowing of the oscillator and may therefore be achieved less efficiently. Finally, the loss of coherent resetting in *timeless* mutants indicates that these responses depend on a functional circadian oscillator rather than solely on acute behavioral responses to light, suggesting that photic information is rapidly incorporated into the molecular clockwork itself.

The dynamics of re-entrainment observed in *Ae aegypti* highlight notable differences from several previously studied insect systems (43–46). Early studies observed that phase advances and delays often required several transient cycles of unstability before stable entrainment was achieved. For example, (43) reported several transient cycles in *Leucophaea maderae* after a 12 h light pulse, while (44) reported similar effects on the eclosion rhythms of *Drosophila pseudoobscura*, after exposure to a 15 min light pulse. More recently, (47)reported that multiple cycles were required before stable resetting is achieved in *Anopheles stephensi* and identified light onset as the critical Zeitgeber, in contrast to our results showing light-offset timing as the main Zeitgeber. However, (48) demonstrated that the duration of transient cycles following a light pulse depended on the circadian time at which the perturbation occurs, with some phases requiring multiple cycles while others showing rapid adjustment, in the flesh fly *Sarcophaga argyrostoma*. This contrasts with our observations, where re-entrainment was achieved rapidly. This is similar to reports by (18) showing re-entrainment within a single cycle in *Aedes aegypti*. Importantly, our findings contrast with previous studies in *Anopheles*. (28) described changes in free-running period following a 1-h light pulse, while the length of the free-running period remained stable across treatments in our assays. This persistence in *Aedes aegypti* supports the presence of a stable core oscillator and suggests that light is a fast-acting entraining signal that resets the phase without modifying the clock’s intrinsic rate.

It is important to note that our experimental design leaves questions that will need to be addressed by future studies. For example, 4 h of light stimulus, whether delivered continuously or in short intermittent bouts, was insufficient to produce a robust phase shift, while 8 h of continuous photic stimulation was. However, the current design did not isolate the causal feature of ON|8 treatment. Indeed, these treatments differed in the amount of light exposure (4 h vs 8 h) and the offset time of the shift (ZT4 vs ZT8) and by extension the deviation between the baseline peak activity time (−8 h vs - 4 h, respectively). This prevents us from concluding about these specific effects. Therefore, future experiments should independently and systematically manipulate the duration and continuity of light exposure and the lights-off timing.

Contrary to *Aedes albopictus, Aedes aegypti* is typically described as endophagic and endophilic (49). Therefore, most bites and transmission events are expected to occur in urban habitats, near or inside human houses that are increasingly equipped with artificial lighting (50). In this context, we designed light cycles with sharp transitions to mimic abrupt changes in indoor light environments. When a mosquito is suddenly confined in a room, both phase-advance and phase-delay scenarios are relevant: the room may remain dark during the day (*e*.*g*., dark rooms, storage areas, garages, basements, windowless rooms), or it may stay illuminated well after sunset due to prolonged human activity in private or public spaces. In both situations, our results show that mosquitoes rapidly adapt to the new light phase and adjust their locomotor activity, becoming more active during the late hours of this new artificial “day”.

A single 1-h light pulse was insufficient to re-entrain circadian locomotor rhythms. Although we did not directly quantify host-seeking or blood feeding under these conditions, previous work in *Anopheles gambiae* suggests that such disruptions can induce marked changes in feeding and transmission-related behaviors (17). These disruptions could be leveraged to complement ongoing efforts to manipulate mosquito behavior using traps and domestic lighting, and highlight the need to better characterize mosquito activity and behavior in urban environments (51, 52). Host-seeking, resting or post-mating behaviors are likely shaped by the light regime within the microhabitat mosquitoes occupy, including the possibility that mosquitoes preferentially exploit shaded zones, or hidden places within the vicinity of human dwellings (53).

Our results also have practical implications for mosquito rearing and experimental design in the laboratory. Many behavioral assays and infection experiments require precise alignment between mosquito peak activity and the experimenter’s time window. In practice, this often involves imposing phase shifts between insectarium and experimental rooms to accommodate experimenters or resource availability. By convention, researchers frequently assume that one to three days of exposure to a new light/dark schedule are sufficient for entrainment (54, 55). Yet, to our knowledge, this is the first study to explicitly quantify the minimal entrainment duration required for mosquitoes to synchronize to a new cycle. Fortunately, *Aedes aegypti* rapidly re-entrains to altered light/dark schedules, and our data indicate that experimental setups should account not only for sufficient entrainment time, but also for the asymmetric effects of phase advances and phase delays when planning time-sensitive assays.

Finally, our work also speaks directly to Sterile Insect Technique (SIT) applications, where a major challenge lies in ensuring that irradiated males retain enough fitness to compete successfully for mates (56). By characterizing how quickly and in which direction mosquito circadian clocks adjust to light shifts, we identify requirements for synchronizing the activity of released males with that of wild females. Optimizing pre-release light regimes could improve mating success and thereby increase the efficacy of SIT programs.

These findings should be interpreted within the broader context of multimodal temporal cues and social interactions. First, extensive literature has examined olfactory signaling in mosquitoes and its role in circadian regulation (57). Heat, sound, and other sensory modalities are also at least partly governed by circadian rhythms, and prior work has shown that olfactory inputs can modulate visual processing in mosquitoes (20, 58, 59). Given the strongly rhythmic nature of olfactory sensitivity in mosquitoes, we hypothesize that phase shifts or disruptions in circadian activity might alter olfactory performance, with downstream consequences for host-seeking, insecticide resistance, sleep, and the insect’s metabolism. These effects are also likely to interact with longer-term photoperiodic effects, such as those associated with seasonal changes or the impact of insecticide-treated bed nets on activity patterns (60, 61).

Finally, in our experiments, we used a standard white light source of approximately 800 lumens, but indoor and outdoor artificial lighting spans a wide range of intensities and spectra, which warrants further investigation. While we focused on *Aedes aegypti* as a diurnal urban mosquito, our simple paradigm provides a comparative framework to explore phase shifts in other vector species and to measure their consequences for host-seeking and blood feeding competence, across diurnal and nocturnal species (19). In this study, we focused our analyses on non-infected females, yet circadian adjustments may also be modulated by social interactions between males and females, as well as by the infection status, given that arboviruses are known to alter mosquito behavior, physiology, and circadian regulation to optimize their own transmission (62, 63).

## Conclusions

Overall, by leveraging the particular ecological context of the urban *Aedes aegypti* mosquito, this work reveals their reliance on a circadian system that is both flexible and highly robust, allowing mosquitoes to immediately adjust their activity patterns to unanticipated changes in the photoperiodic cycles. These properties have implications for urban vector populations and open new perspectives for using light as an innovative vector control strategy. In the context of a continuously-growing human population, a rapid urbanization, and an expanding use of light-emitting electronic devices, it becomes highly significant to better understand how disease vectors adapt to these changes so we can identify intervention points to reduce or redirect their vectorial capacity.

## ACKNOWLEDGEMENTS

The authors thank Danny Eanes for technical assistance, and Seyed Jalil Pasha (JP) Mirlohi and Shajaesza Diggs for assistance with maintaining the mosquito lines used in this study. Dr. Chloé Lahondère for fruitful discussions and feedback. The following reagents were obtained through BEI Resources, NIAID, NIH: *Aedes aegypti*, Strain LVP-IB12, Eggs, MRA-735, and Strain ROCK, MRA-734, both contributed by David W. Severson. This work received direct support from the National Institute of Allergy and Infectious Diseases (NIAID) of the National Institutes of Health under Award R01AI155785, which investigates the rhythms in mosquito olfactory processes, as well as from the National Institute of Food and Agriculture, U.S. Department of Agriculture, through Hatch project VA-160212 (to C.V.).

## Supplementary Figures

**Fig S1.**
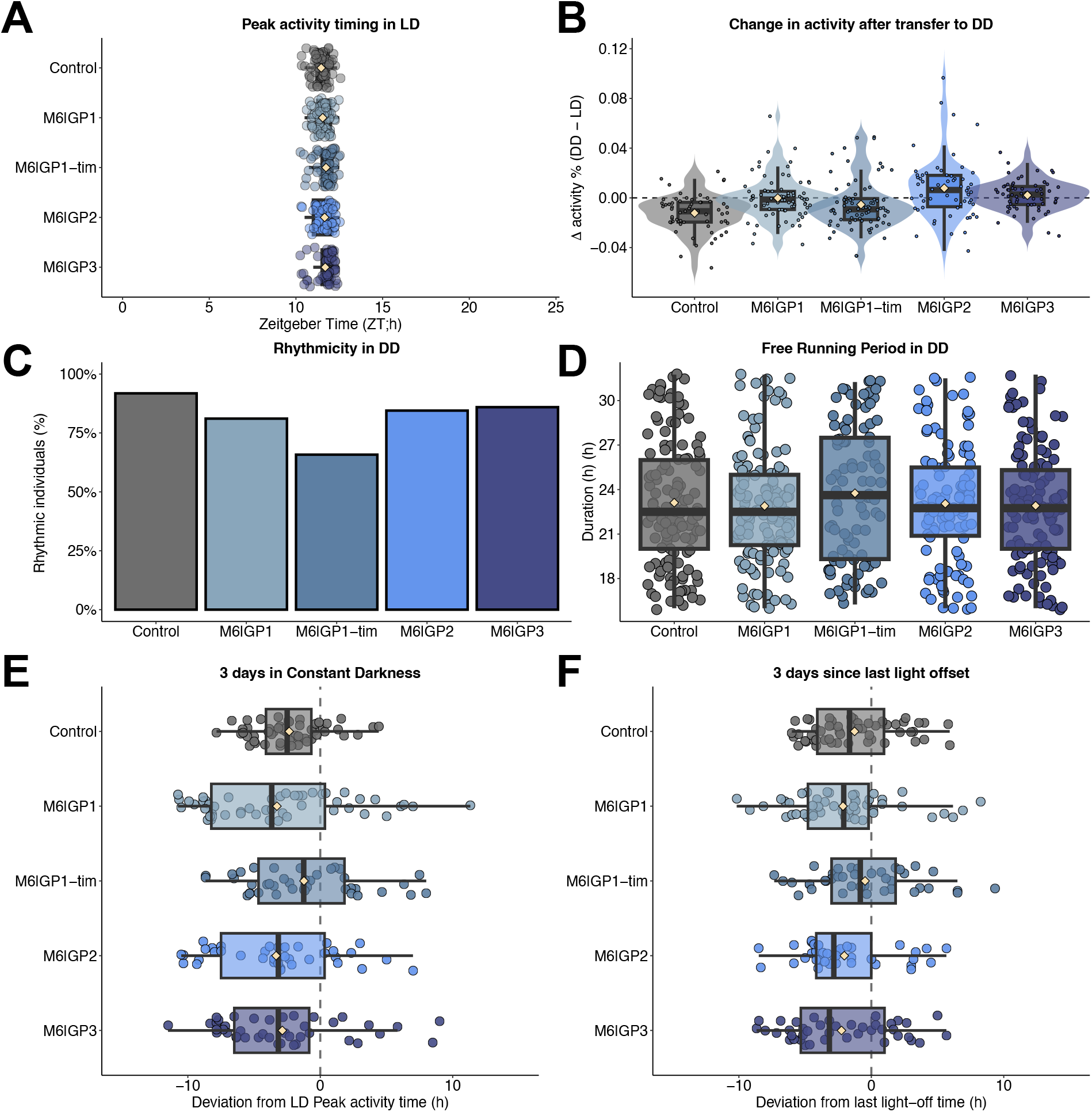
Population profile of mosquitoes following a 6-h phase advance and a 24-h recovery period. **(A)** Peak activity timing in LD conditions. Each point represents an individual’s activity peak phase. **(B)** Mean activity change from LD to DD conditions. Violin plots show the distribution of individual values, and points represent individual mosquitoes. **(C-D)** Rhythmicity in DD. Locomotor activity period was estimated for each individual using an autocorrelation periodogram. For each individual, the highest significant peak per 24-h period was retained for the three days in DD. **(B)** Bars represent the proportion of rhythmic individuals in each group. **(C)** Points represent individual period estimates. **(E-F)** 3-days average of phase deviation of activity peaks timing relative to **(E)** constant darkness transition and **(F)** the last light-off event, similar to Fig. 1 to 5 in the main text, calculated on the three days in DD instead of the first 24-h of DD. Each point represents an individual’s activity peak phase. For all: boxes show the interquartile range and median, whiskers indicate the distribution spread, points represent individual observations, and beige diamond-shaped symbols indicate group means.

**Fig S2.**
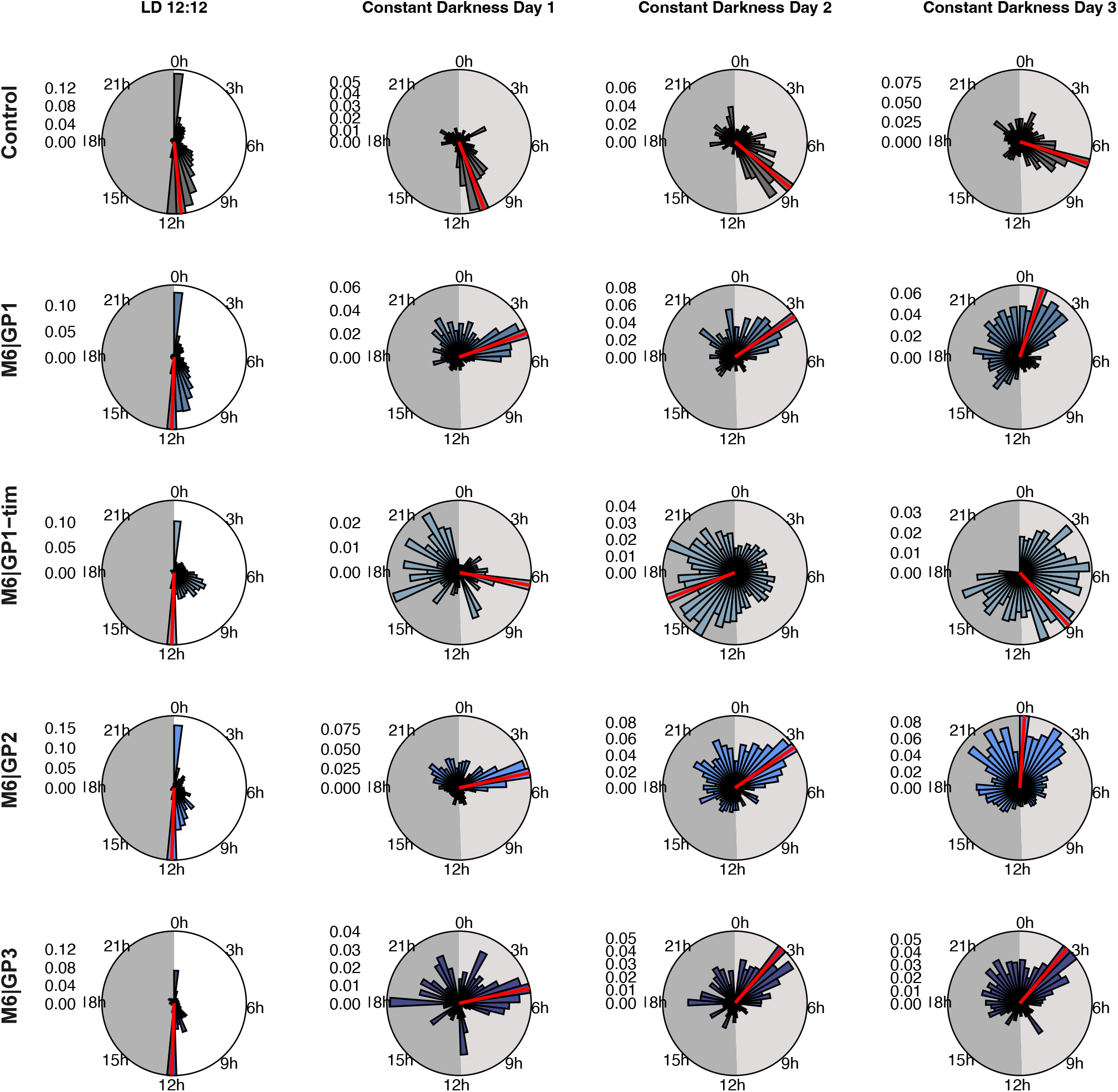
Daily activity of mosquitoes following a 6-h phase advance and a 24-h recovery period. Rose diagrams show the mean activity calculated in 30-min bins across 24 h. Panels show the LD 12:12 cycle followed by days 1, 2 and 3 in constant darkness (DD1-DD3), from left to right. Under LD conditions, the shaded sector denotes the dark phase. Under DD conditions, the light- and dark-grey sectors indicate the subjective day and subjective night, respectively, based on the preceding LD cycle. Each panel is normalized to its own maximum activity. The black line indicates the maximum mean activity for each group, while the red radial line marks the time bin containing the activity peak.

**Fig S3.**
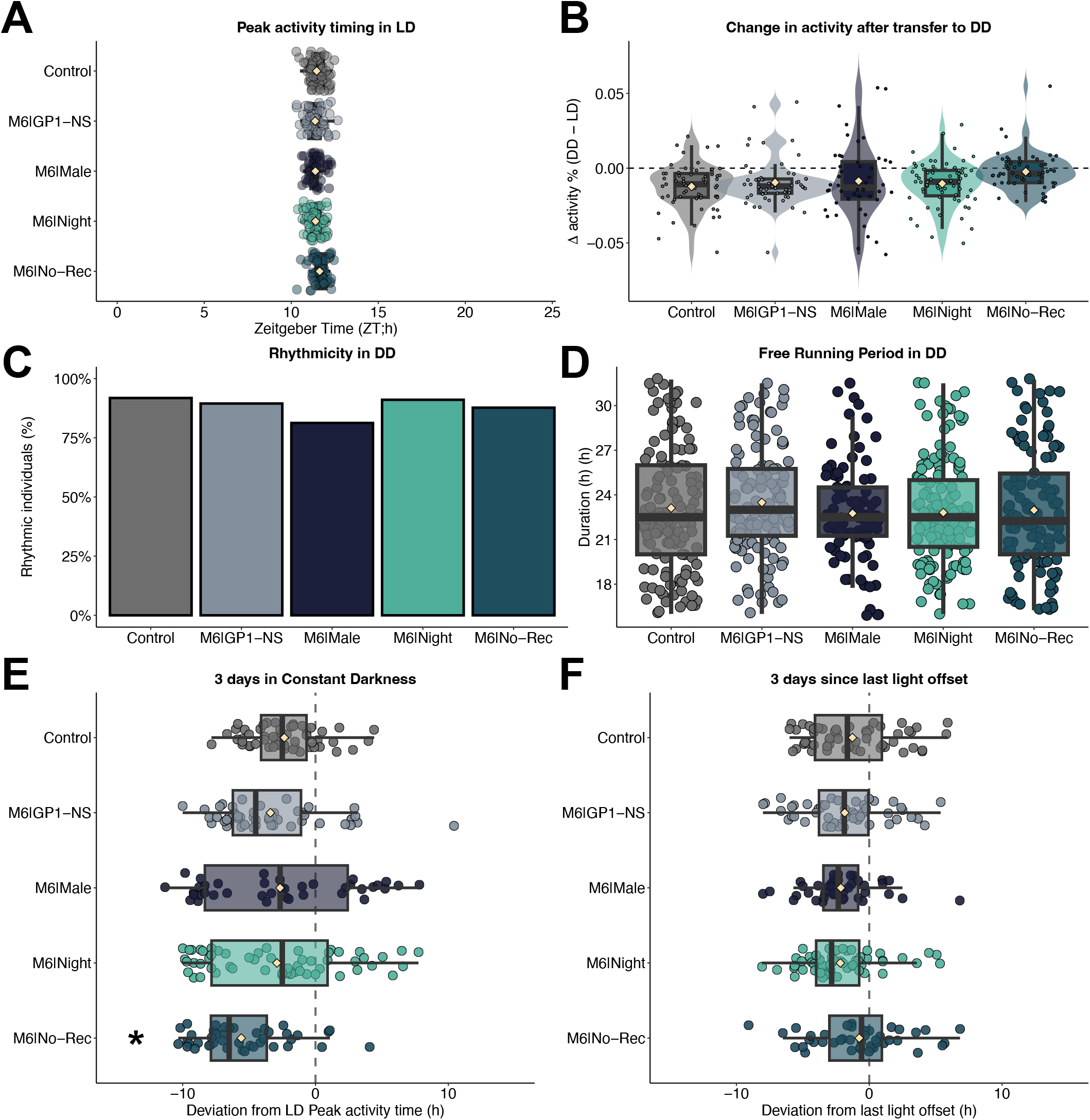
Population profile of mosquitoes following a 6-h phase advance reserved, for males, during the scotophase or with no recovery period. **(A)** Peak activity timing in LD conditions. Each point represents an individual’s activity peak phase. **(B)** Mean activity change from LD to DD conditions. Violin plots show the distribution of individual values, and points represent individual mosquitoes. **(C-D)** Rhythmicity in DD. Locomotor activity period was estimated for each individual using an autocorrelation periodogram. For each individual, the highest significant peak per 24-h period was retained for the three days in DD. **(B)** Bars represent the proportion of rhythmic individuals in each group. **(C)** Points represent individual period estimates. **(E-F)** 3-days average of phase deviation of activity peaks timing relative to **(E)** constant darkness transition and **(F)** the last light-off event, similar to Fig. 1 to 5 in the main text, calculated on the three days in DD instead of the first 24-h of DD. Each point represents an individual’s activity peak phase. For all: boxes show the interquartile range and median, whiskers indicate the distribution spread, points represent individual observations, and beige diamond-shaped symbols indicate group means.

**Fig S4.**
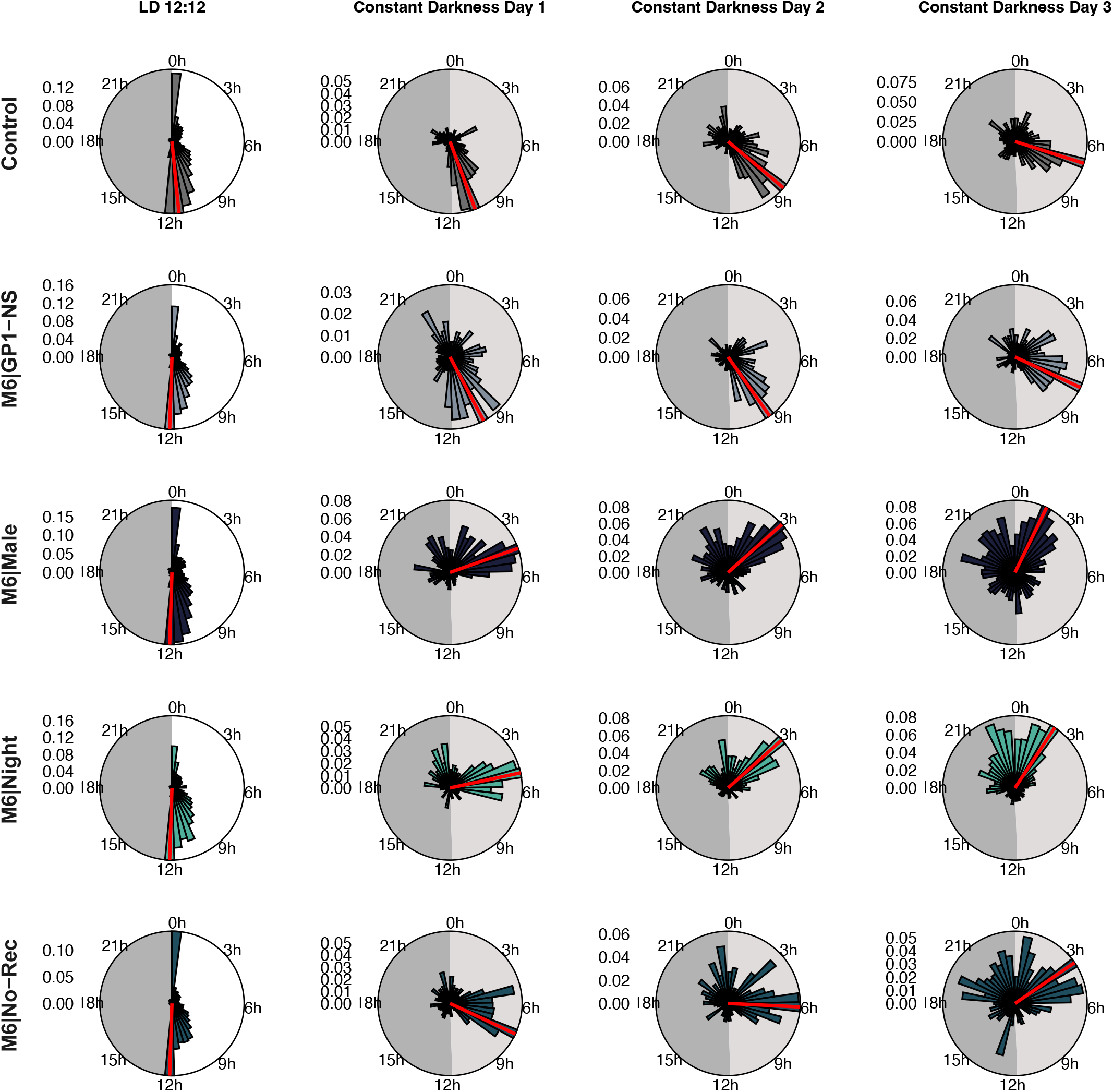
Daily activity of mosquitoes following a 6-h phase advance reserved, for males, during the scotophase or with no recovery period. Rose diagrams show the mean activity calculated in 30-min bins across 24 h. Panels show the LD 12:12 cycle followed by days 1, 2 and 3 in constant darkness (DD1-DD3), from left to right. Under LD conditions, the shaded sector denotes the dark phase. Under DD conditions, the light- and dark-grey sectors indicate the subjective day and subjective night, respectively, based on the preceding LD cycle. Each panel is normalized to its own maximum activity. The black line indicates the maximum mean activity for each group, while the red radial line marks the time bin containing the activity peak.

**Fig S5.**
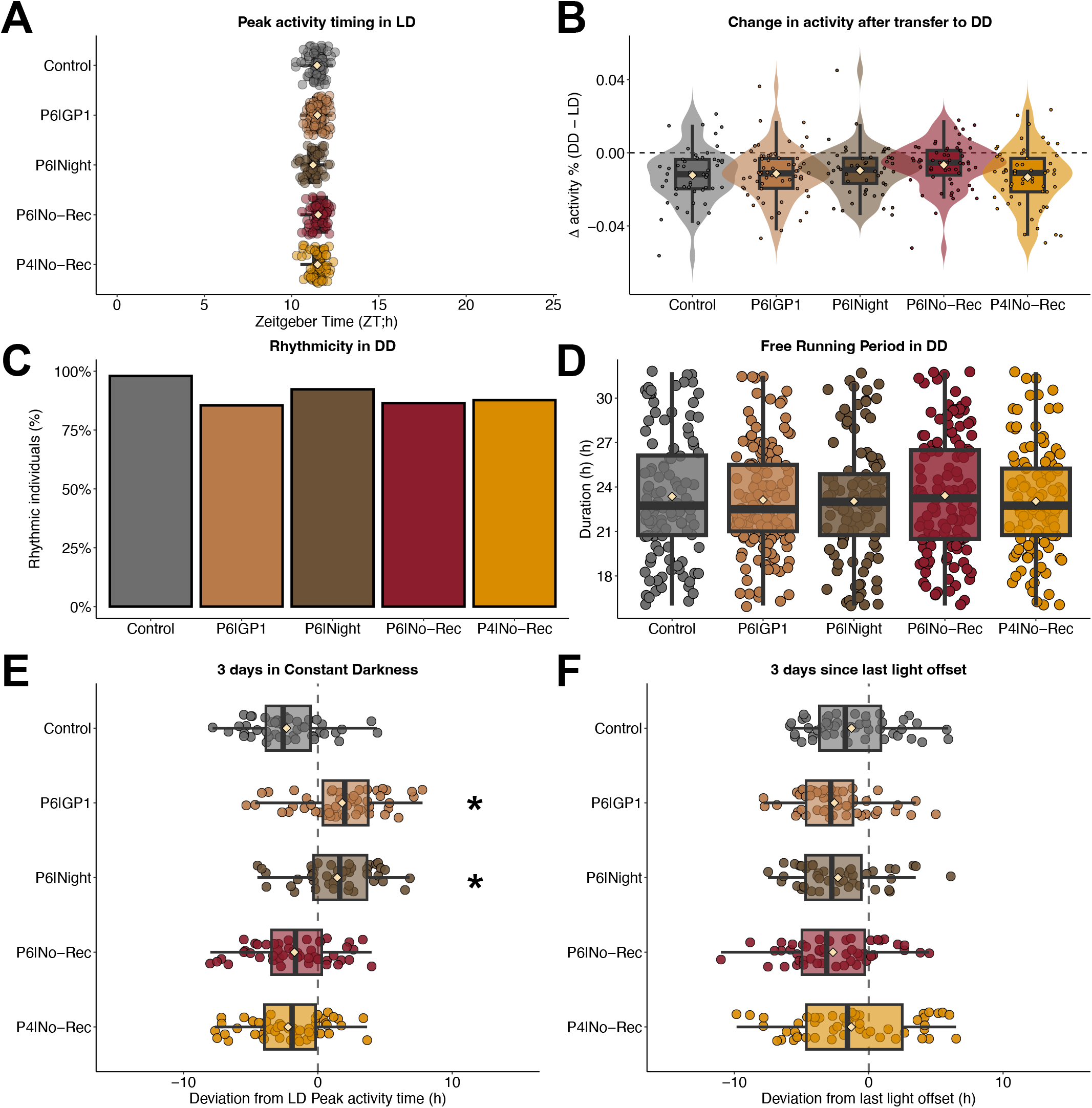
Population profile of mosquitoes following a 6-h phase delay. **(A)** Peak activity timing in LD conditions. Each point represents an individual’s activity peak phase. **(B)** Mean activity change from LD to DD conditions. Violin plots show the distribution of individual values, and points represent individual mosquitoes. **(C-D)** Rhythmicity in DD. Locomotor activity period was estimated for each individual using an autocorrelation periodogram. For each individual, the highest significant peak per 24-h period was retained for the three days in DD. **(B)** Bars represent the proportion of rhythmic individuals in each group. **(C)** Points represent individual period estimates. **(E-F)** 3-days average of phase deviation of activity peaks timing relative to **(E)** constant darkness transition and **(F)** the last light-off event, similar to Fig. 1 to 5 in the main text, calculated on the three days in DD instead of the first 24-h of DD. Each point represents an individual’s activity peak phase. For all: boxes show the interquartile range and median, whiskers indicate the distribution spread, points represent individual observations, and beige diamond-shaped symbols indicate group means.

**Fig S6.**
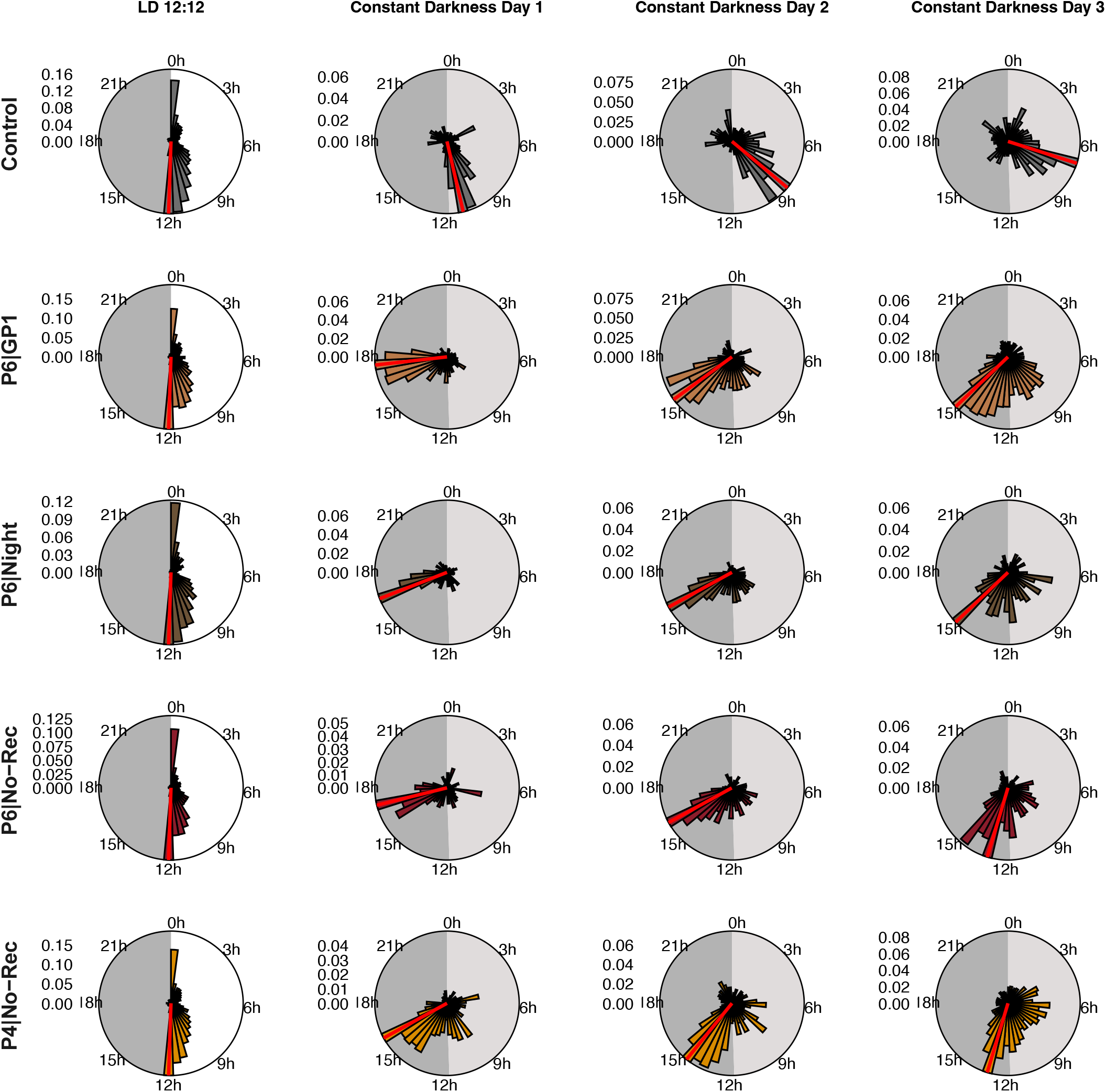
Daily activity of mosquitoes following a 6-h phase delay. Rose diagrams show the mean activity calculated in 30-min bins across 24 h. Panels show the LD 12:12 cycle followed by days 1, 2 and 3 in constant darkness (DD1-DD3), from left to right. Under LD conditions, the shaded sector denotes the dark phase. Under DD conditions, the light- and dark-grey sectors indicate the subjective day and subjective night, respectively, based on the preceding LD cycle. Each panel is normalized to its own maximum activity. The black line indicates the maximum mean activity for each group, while the red radial line marks the time bin containing the activity peak.

**Fig S7.**
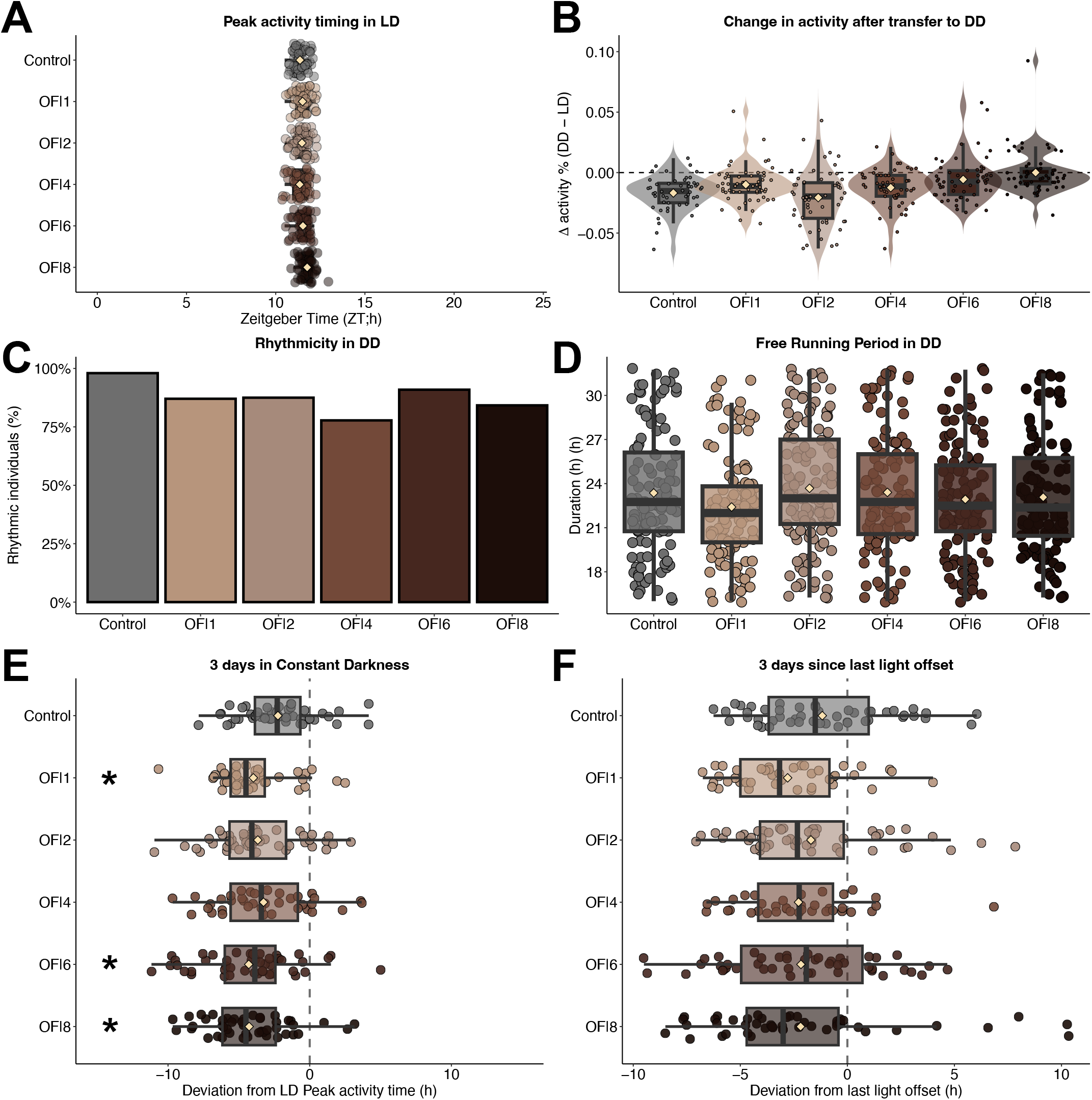
Population profile of mosquitoes following a 6-h phase advance aligned with the lights-off. **(A)** Peak activity timing in LD conditions. Each point represents an individual’s activity peak phase. **(B)** Mean activity change from LD to DD conditions. Violin plots show the distribution of individual values, and points represent individual mosquitoes. **(C-D)** Rhythmicity in DD. Locomotor activity period was estimated for each individual using an autocorrelation periodogram. For each individual, the highest significant peak per 24-h period was retained for the three days in DD. **(B)** Bars represent the proportion of rhythmic individuals in each group. **(C)** Points represent individual period estimates. **(E-F)** 3-days average of phase deviation of activity peaks timing relative to **(E)** constant darkness transition and **(F)** the last light-off event, similar to Fig. 1 to 5 in the main text, calculated on the three days in DD instead of the first 24-h of DD. Each point represents an individual’s activity peak phase. For all: boxes show the interquartile range and median, whiskers indicate the distribution spread, points represent individual observations, and beige diamond-shaped symbols indicate group means.

**Fig S8.**
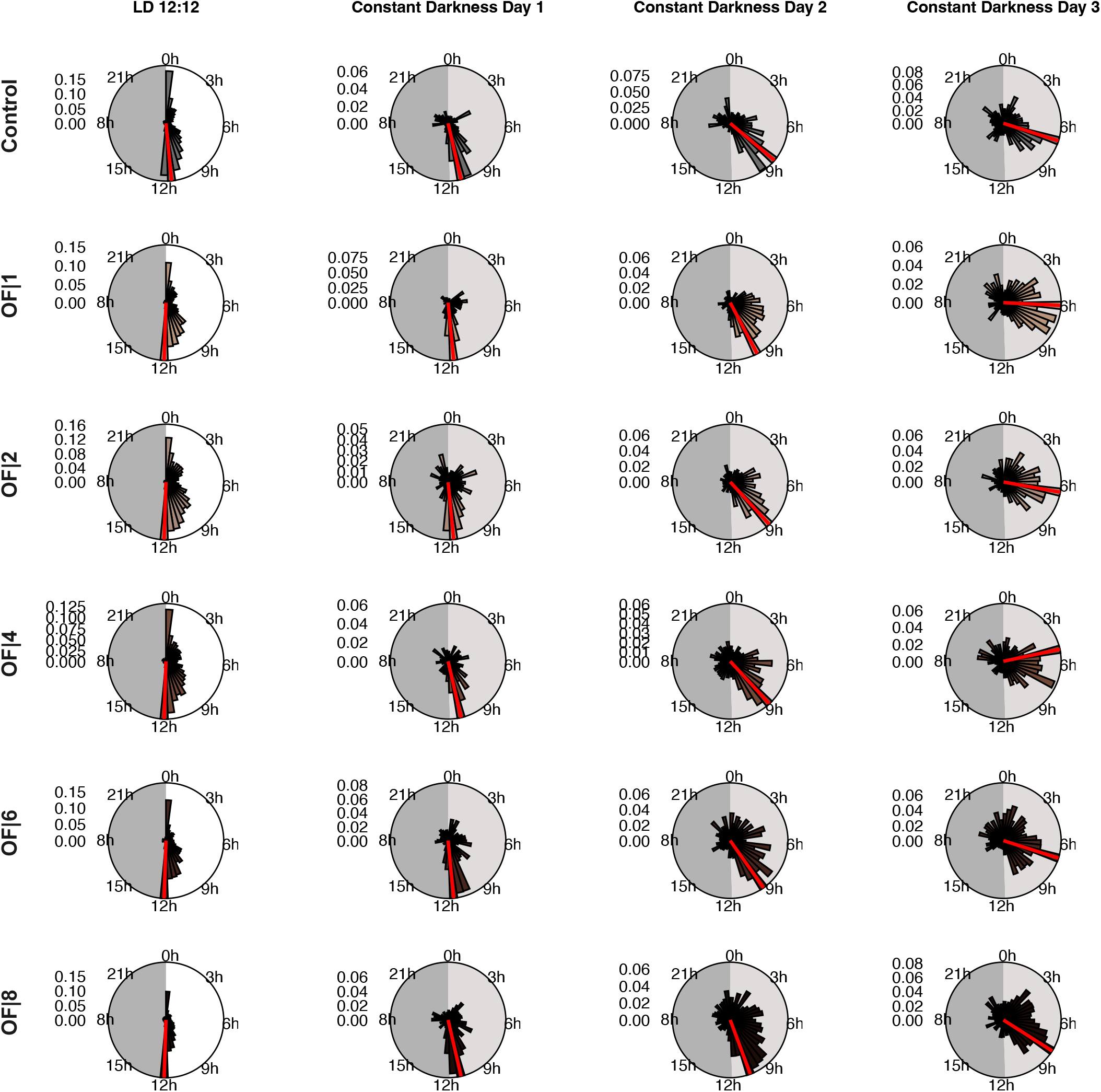
Daily activity of mosquitoes following a 6-h phase advance aligned with the lights-off. Rose diagrams show the mean activity calculated in 30-min bins across 24 h. Panels show the LD 12:12 cycle followed by days 1, 2 and 3 in constant darkness (DD1-DD3), from left to right. Under LD conditions, the shaded sector denotes the dark phase. Under DD conditions, the light- and dark-grey sectors indicate the subjective day and subjective night, respectively, based on the preceding LD cycle. Each panel is normalized to its own maximum activity. The black line indicates the maximum mean activity for each group, while the red radial line marks the time bin containing the activity peak.

**Fig S9.**
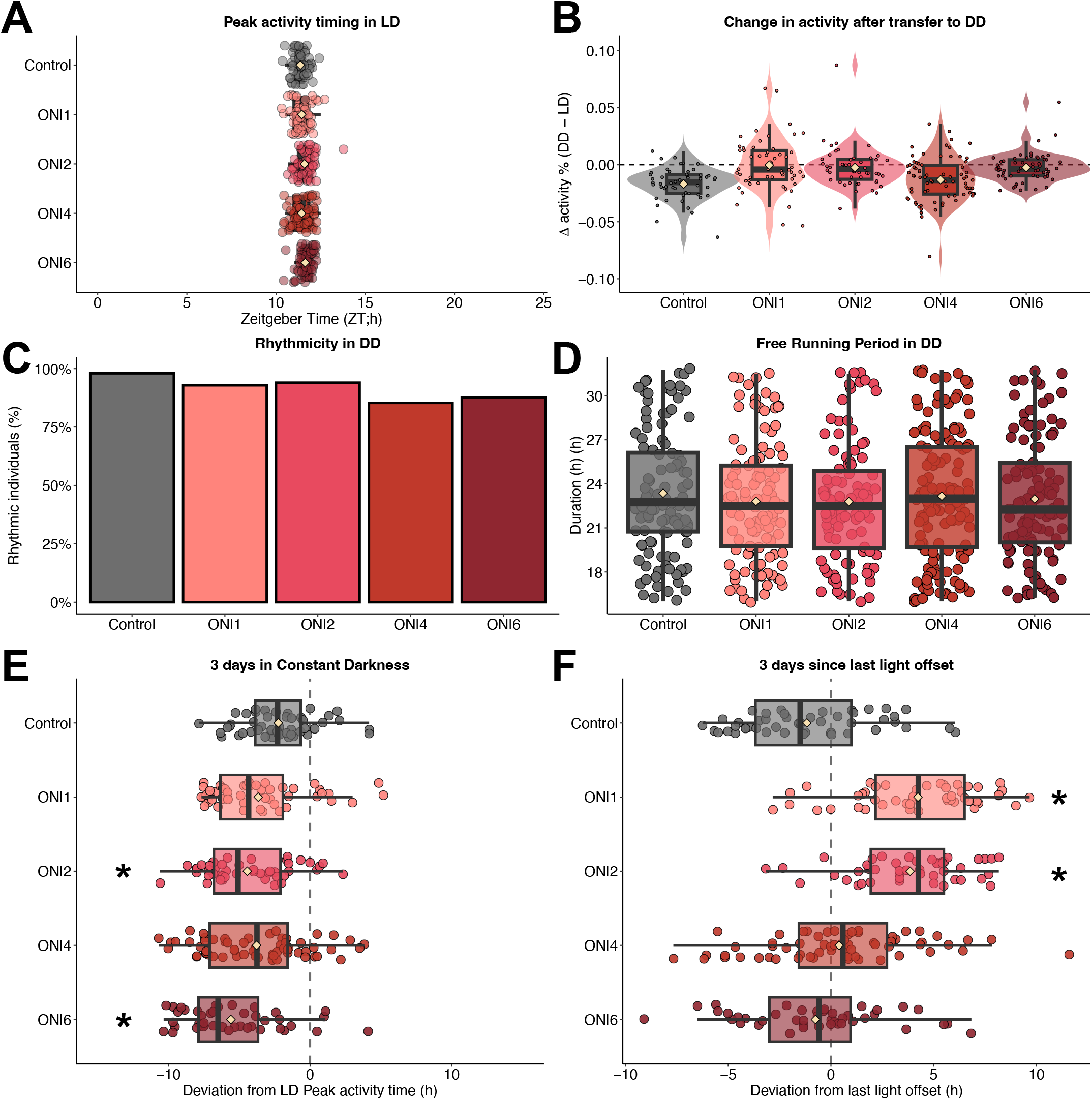
Population profile of mosquitoes following a 6-h phase advance aligned with the lights-on. **(A)** Peak activity timing in LD conditions. Each point represents an individual’s activity peak phase. ON|6 treatment corresponds to M6|No-Rec treatment in (Fig. 2). **(B)** Mean activity change from LD to DD conditions. Violin plots show the distribution of individual values, and points represent individual mosquitoes. **(C-D)** Rhythmicity in DD. Locomotor activity period was estimated for each individual using an autocorrelation periodogram. For each individual, the highest significant peak per 24-h period was retained for the three days in DD. **(B)** Bars represent the proportion of rhythmic individuals in each group. **(C)** Points represent individual period estimates. **(E-F)** 3-days average of phase deviation of activity peaks timing relative to **(E)** constant darkness transition and **(F)** the last light-off event, similar to Fig. 1 to 5 in the main text, calculated on the three days in DD instead of the first 24-h of DD. Each point represents an individual’s activity peak phase. For all: boxes show the interquartile range and median, whiskers indicate the distribution spread, points represent individual observations, and beige diamond-shaped symbols indicate group means.

**Fig S10.**
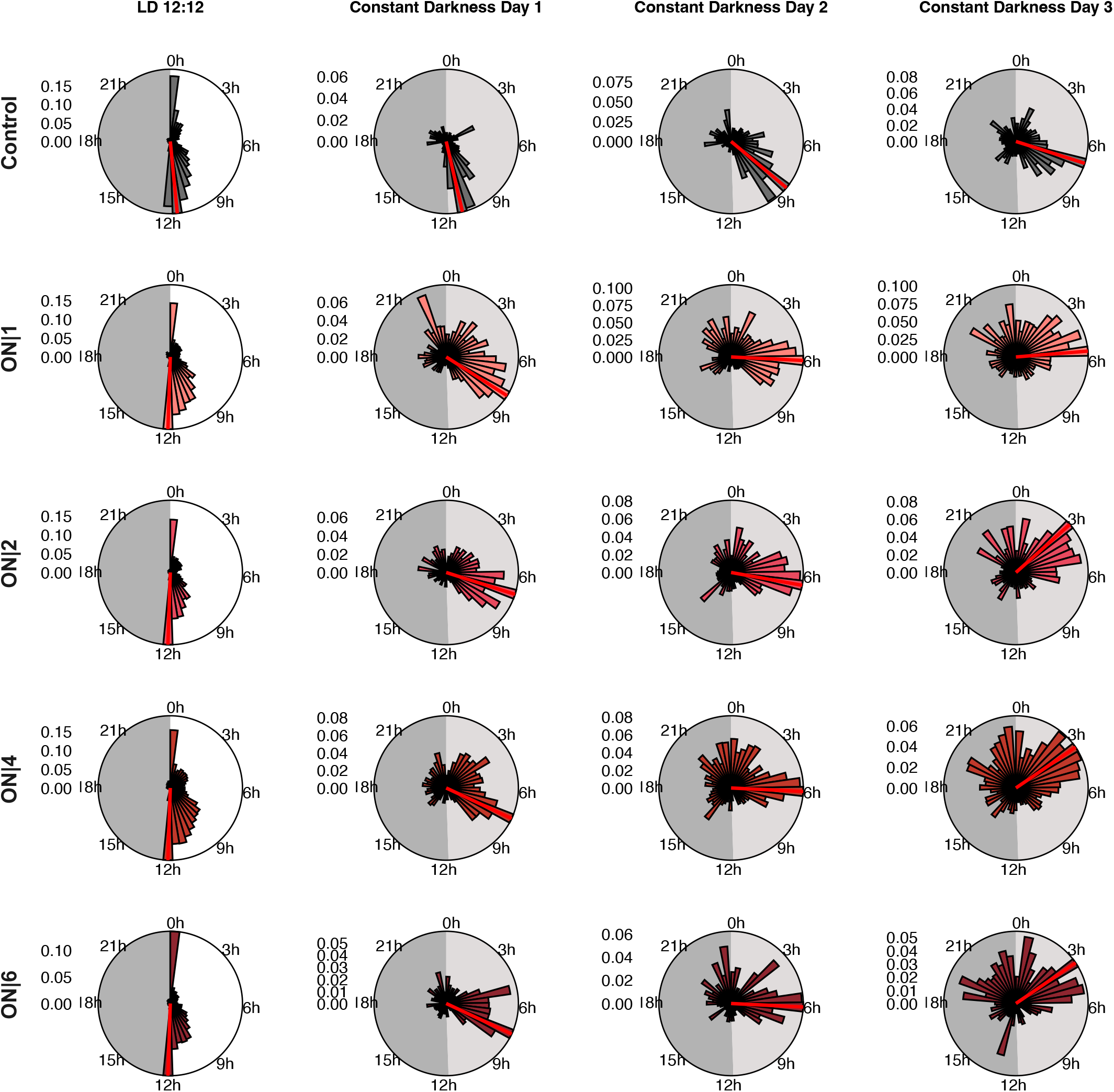
Daily activity of mosquitoes following a 6-h phase advance aligned with the lights-on. Rose diagrams show the mean activity calculated in 30-min bins across 24 h. Panels show the LD 12:12 cycle followed by days 1, 2 and 3 in constant darkness (DD1- DD3), from left to right. Under LD conditions, the shaded sector denotes the dark phase. Under DD conditions, the light- and dark-grey sectors indicate the subjective day and subjective night, respectively, based on the preceding LD cycle. Each panel is normalized to its own maximum activity. The black line indicates the maximum mean activity for each group, while the red radial line marks the time bin containing the activity peak.

**Fig S11.**
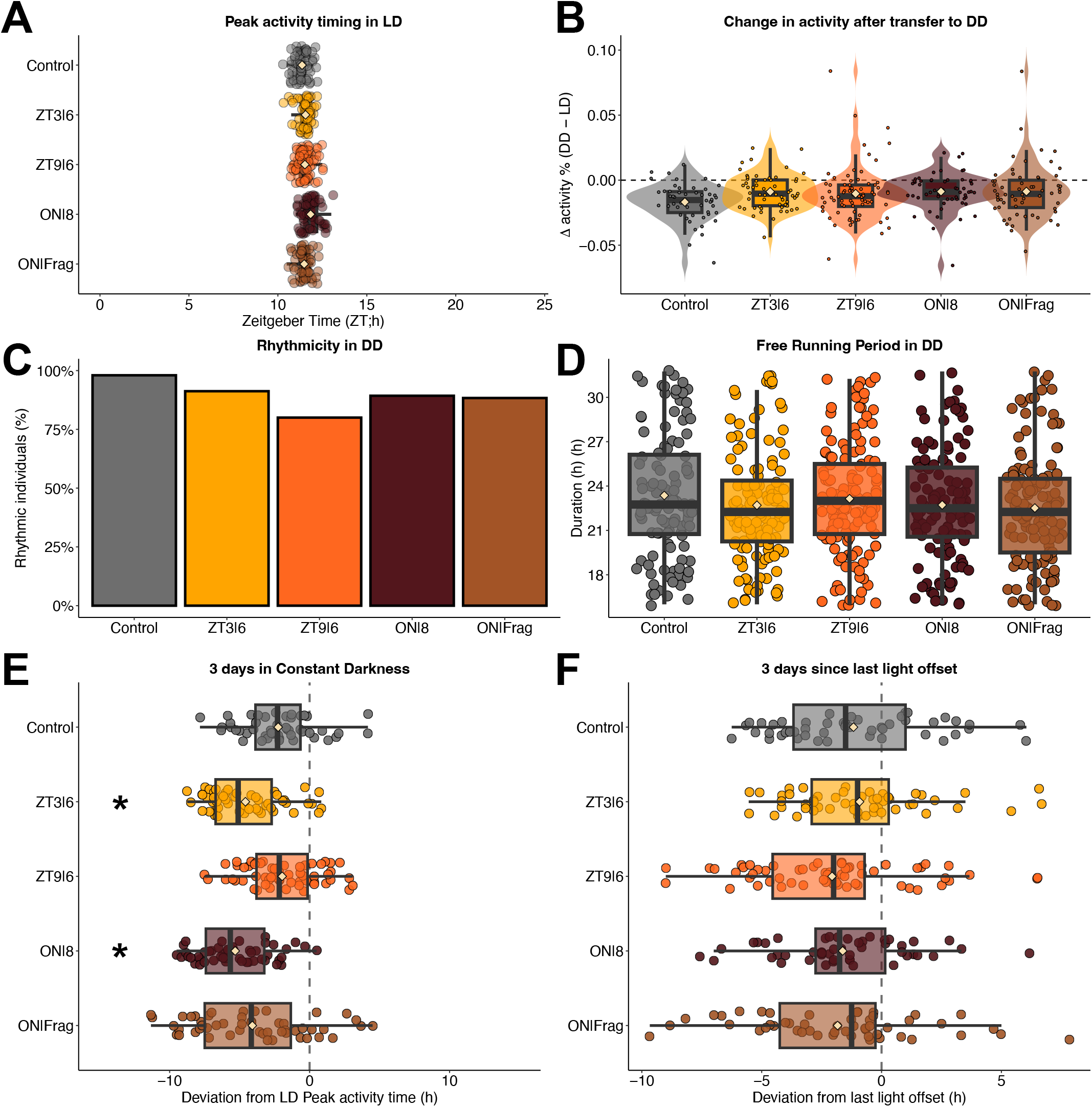
Population profile of mosquitoes following a 6 or 8-h phase advance aligned at different time, and with continuous or fragmented light exposure. **(A)** Peak activity timing in LD conditions. Each point represents an individual’s activity peak phase.**(A)** Mean activity change from LD to DD conditions. Violin plots show the distribution of individual values, and points represent individual mosquitoes. **(C-D)** Rhythmicity in DD. Locomotor activity period was estimated for each individual using an autocorrelation periodogram. For each individual, the highest significant peak per 24-h period was retained for the three days in DD. **(B)** Bars represent the proportion of rhythmic individuals in each group. **(C)** Points represent individual period estimates. **(E-F)** 3-days average of phase deviation of activity peaks timing relative to **(E)** constant darkness transition and **(F)** the last light-off event, similar to Fig. 1 to 5 in the main text, calculated on the three days in DD instead of the first 24-h of DD. Each point represents an individual’s activity peak phase. For all: boxes show the interquartile range and median, whiskers indicate the distribution spread, points represent individual observations, and beige diamond-shaped symbols indicate group means.

**Fig S12.**
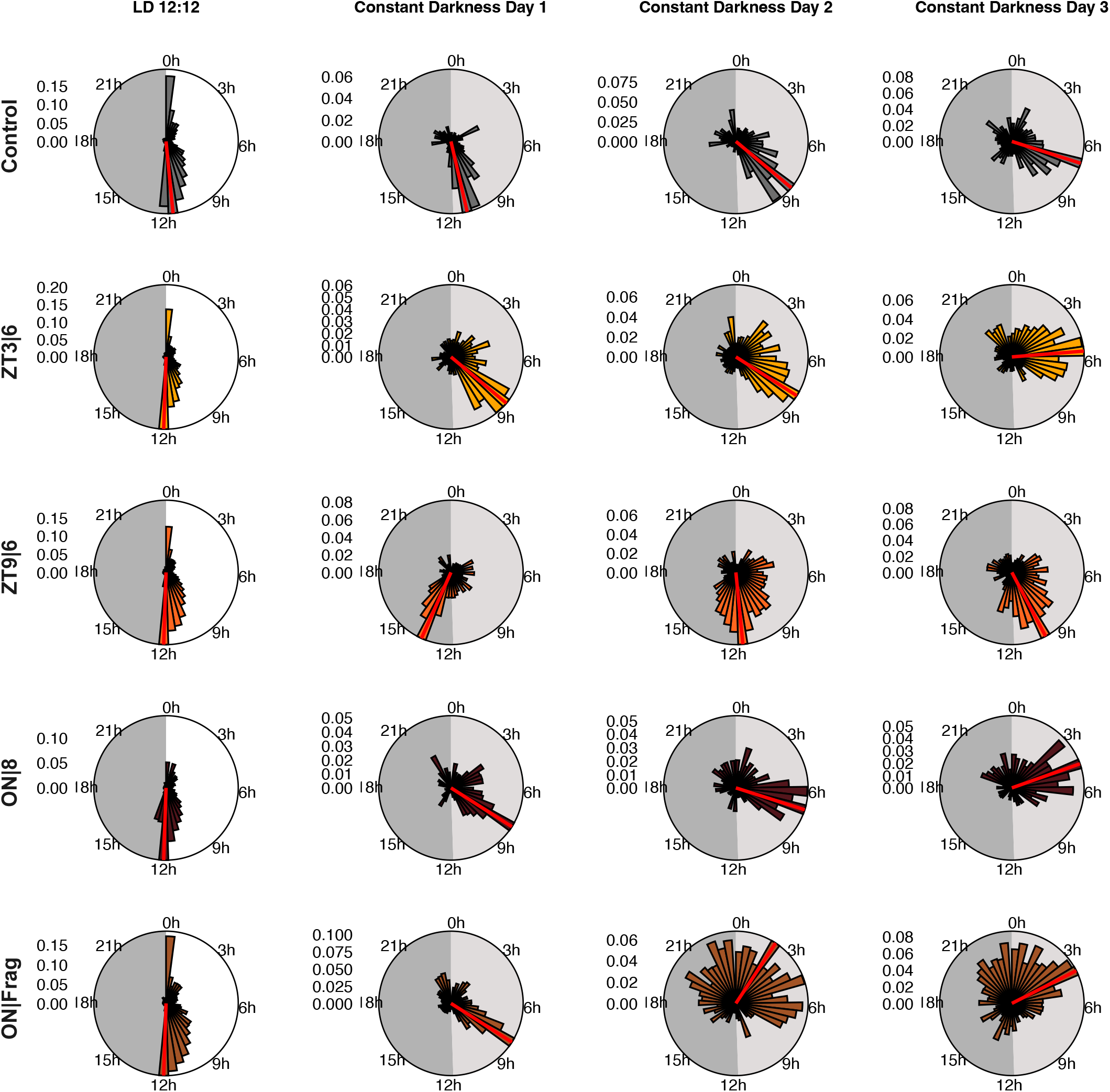
Daily activity of mosquitoes following a 6 or 8-h phase advance aligned at different time, and with continuous or fragmented light exposure. Rose diagrams show the mean activity calculated in 30-min bins across 24 h. Panels show the LD 12:12 cycle followed by days 1, 2 and 3 in constant darkness (DD1-DD3), from left to right. Under LD conditions, the shaded sector denotes the dark phase. Under DD conditions, the light- and dark-grey sectors indicate the subjective day and subjective night, respectively, based on the preceding LD cycle. Each panel is normalized to its own maximum activity. The black line indicates the maximum mean activity for each group, while the red radial line marks the time bin containing the activity peak.

## Notes

### Competing Interest Statement

The authors have declared no competing interest.

